# Integrative phenotypic-transcriptomic analysis of soybean plants subjected to multifactorial stress combination

**DOI:** 10.1101/2023.08.15.553447

**Authors:** María Ángeles Peláez-Vico, Ranjita Sinha, Sai Preethi Induri, Zhen Lyu, Sai Darahas Venigalla, Dinesh Vasireddy, Pallav Singh, Manish Sridhar Immadi, Lidia S. Pascual, Benjamin Shostak, David Mendoza-Cózatl, Trupti Joshi, Felix B. Fritschi, Sara I. Zandalinas, Ron Mittler

**Author notes:** Author for correspondence and distribution of materials: Ron Mittler. These authors contributed equally (Ranjita Sinha, Sai Preethi Induri).

## Abstract

Global warming, climate change, and industrial pollution are altering our environment subjecting crops to an increasing number and complexity of abiotic stress conditions, concurrently or sequentially. Recent studies revealed that a combination of 3 or more stresses simultaneously impacting a plant (termed ‘multifactorial stress combination’; MFSC) can cause a drastic decline in plant growth and survival, even if the level of each stress involved in the MFSC has a negligible effect on plants. However, the impacts of MFSC on crops are largely unknown. We subjected soybean plants to a MFSC of up to five different stresses (water deficit, salinity, low phosphate, acidity, and cadmium), in an increasing level of complexity, and conducted integrative transcriptomic-phenotypic analysis of reproductive and vegetative tissues. We reveal that MFSC has a negative cumulative effect on soybean yield, that each set of MFSC condition elicits a unique transcriptomic response (that is different between flowers and leaves), and that selected genes expressed in leaves or flowers are linked to the effects of MFSC on different vegetative, physiological, and/or reproductive parameters. We further reveal that the transcriptomic response of soybean and *Arabidopsis* to MFSC shares common features associated with reactive oxygen and iron/copper signaling/metabolism. Our study provides unique phenotypic and transcriptomic datasets for dissecting the mechanistic effects of MFSC on the vegetative, physiological, and reproductive processes of a crop plant.

## INTRODUCTION

Global warming and climate change are altering our environment causing drastic changes in weather patterns at different parts of the world (Mazdiyasni and AghaKouchak, 2015; Alizadeh et al., 2020; Masson-Delmotte et al., 2021; Rivero et al., 2022). This process is subjecting crops to new types of abiotic stresses, not common to these areas, as well as to heightened intensities of stresses normal to them (Long and Ort, 2010; Mittler and Blumwald, 2010; Bailey-Serres et al., 2019; Zandalinas et al., 2021a). The most common types of abiotic stresses that impact crops are drought, flooding, heat waves, salinity, nutrient deprivation, and cold/freezing stress, as well as their simultaneous and/or consecutive occurrences, that may happen during the vegetative and/or reproductive phases of the crop growth cycle (Mooney et al., 1991; Nilsen and Orcutt, 1996; Mittler, 2006; Prasch and Sonnewald, 2013; Coolen et al., 2016; Shaar-Moshe et al., 2017; Zandalinas et al., 2020a, 2020b; Zsögön et al., 2021). These abiotic stresses, and their combinations, cause decreases in crop yield and destabilize the security and economy of different countries around the globe (Mittler and Blumwald, 2010; Lobell et al., 2011; Challinor et al., 2014; Bailey-Serres et al., 2019). Accompanying these abiotic stresses are often changes in the population dynamics and distribution of pathogens, insects, pests, weeds, and/or different invasive species, that could also have a negative impact on crop yield (Huang et al., 2020; Hamann et al., 2021). Adding to the potential negative impacts of the possible combinations of abiotic and/or biotic stresses described above, is the overall increase in environmental pollutions (other than greenhouse gasses), caused by human activity, that could also negatively impact crop yield (Masson-Delmotte et al., 2021; Maddela et al., 2022). Examples of these include different chemicals that pollute our air, water, and/or soil, such as persistent organic compounds, microplastics, ozone, heavy metals, pesticides, and/or herbicides, that can also negatively impact crop growth and yield (Zandalinas et al., 2021a; Shrestha et al., 2022). Taken together, the growing, mostly unabated, and complex effects of human activity on our natural environment could potentially subject crops, soils, and microbiomes to different combinations of multiple stresses, simultaneously or sequentially (Côté et al., 2016; Sage, 2020; Zhou et al., 2020; Rillig et al., 2021a; Zandalinas et al., 2021a). We and others have recently termed this effect ‘multifactorial stress combination’ (MFSC) and begun studying it as a new type of stress that could potentially impact crop yield worldwide (Rillig et al., 2019, 2021b; Zandalinas et al., 2021a, 2021b; Zandalinas and Mittler, 2022; Yang et al., 2022; Pascual et al., 2023; Sinha et al., 2022a). In our first direct study of MFSC effects on plants (using *Arabidopsis thaliana* seedlings), we demonstrated that the effects of a MFSC of up to six different stresses (each applied at a low level that had negligible effects on plants), applied in an increasing complexity to plants, had an overall cumulative negative impact on plant growth and survival (Zandalinas et al., 2021b). This finding revealed a new principle in plant stress biology, termed the ‘MFSC principle’ (Zandalinas et al., 2021a; Mittler and Zandalinas, 2022). This principle states that with the increase in the number and complexity of multiple stresses affecting a plant, plant growth and survival will decline, even if the effects of each stress involved in the MFSC on plants are minimal (Zandalinas et al., 2021a; Mittler and Zandalinas, 2022). We also demonstrated that each different MFSC condition had a unique transcriptomic footprint and that the response of plants to MFSC was altered in Arabidopsis mutants deficient in reactive oxygen species (ROS) signaling and scavenging (Zandalinas et al., 2021b). In two recent studies, we also revealed that the impacts of MFSC on crop plants such as rice (*Oryza sativa*), maize (*Zea mays*), and tomato (*Solanum lycopersicum*), were similar to that on Arabidopsis, in terms of growth and biomass impairment (Pascual et al., 2023; Sinha et al., 2022a), and that MFSC had a cumulative effect on different physiological aspects of tomato, that were altered in a jasmonic acid (JA) signaling mutant (Pascual et al., 2023). We further reported that the proteomics response of rice plants to each MFSC condition was unique (Sinha et al., 2022a). However, the impacts of MFSC on the yield and transcriptomic responses of flowers and leaves of a crop plant were not previously reported. These responses are critical to our understanding of MFSC impacts on yield that could be linked to vegetative and/or reproductive processes, as well as to the molecular and physiological responses of these tissues to MFSC. In this respect, it is important to note that the vegetative and reproductive transcriptomic responses of the crop plant soybean (*Glycine max*) were recently demonstrated to be very different from each other in plants subjected to drought, heat stress, or a combination of drought and heat stress (Sinha et al., 2022b, 2023a, 2023b).

To begin dissecting the effects of MFSC on crop yield and to provide valuable transcriptomic and phenotypic datasets of reproductive and vegetative tissues of a crop plant to MFSC, we subjected soybean plants to a MFSC of up to five different stresses (water deficit, salinity, low phosphate, acidity, and the heavy metal cadmium), in an increasing level of complexity, and studied the impacts of MFSC on vegetative growth, plant physiology, reproductive parameters, and transcriptomic responses of leaves and flowers. Here, we provide unique phenotypic and transcriptomic datasets of a crop plant subjected to MFSC. Using various types of data analyses, we further demonstrate that each set of MFSC conditions elicits a unique transcriptomic response (that is different between flowers and leaves), that the impacts of MFSC on soybean reproductive success are cumulative (overall, following the MFSC principle), and that selected genes expressed in leaves or flowers of soybean during MFSC could be linked to the effects of MFSC on different vegetative, physiological, and reproductive parameters. Our study provides integrative phenotypic-transcriptomic datasets for dissecting the effects of MFSC on the vegetative, physiological, and reproductive processes of a crop plant.

## RESULTS

### The impact of MFSC on soybean growth and biomass

To study the effects of MFSC on a crop plant, we subjected soybean (*Glycine max*) plants growing in a greenhouse to a MFSC of water deficit, salinity, low phosphate, and acidity in all possible combinations, as well as added cadmium stress as a fifth stress to the combination of all four stresses (Figure 1A-1C; similar to the design used for the study of MFSC in Arabidopsis; Zandalinas et al., 2021b). We then measured different growth, physiological, and reproductive parameters and sampled leaves and whole flowers for transcriptomic analysis (Figure 1B, 1C). As shown in Figures 1D-1F, with the increase in the complexity and number of stresses applied to soybean plants, plant growth rate, height, and biomass declined (see also Supplemental Figure S1). In contrast, changes in relative water content (RWC) and total chlorophyll content of plants were not similarly impacted by the MFSC, and only the combined effect of all 5 stresses had a significant negative impact on RWC and total chlorophyll content (Figure 1G, 1H; Supplemental Figure S2). Taken together, the results shown in Figure 1 reveal that although each stress applied individually had a marginal or low impact on plants, the cumulative impact of the different stresses on plant growth rate, height, and biomass were significantly more severe, largely following the basic principle of MFSC in plants (Zandalinas et al., 2021a; Mittler and Zandalinas, 2022).

**Figure 1.**
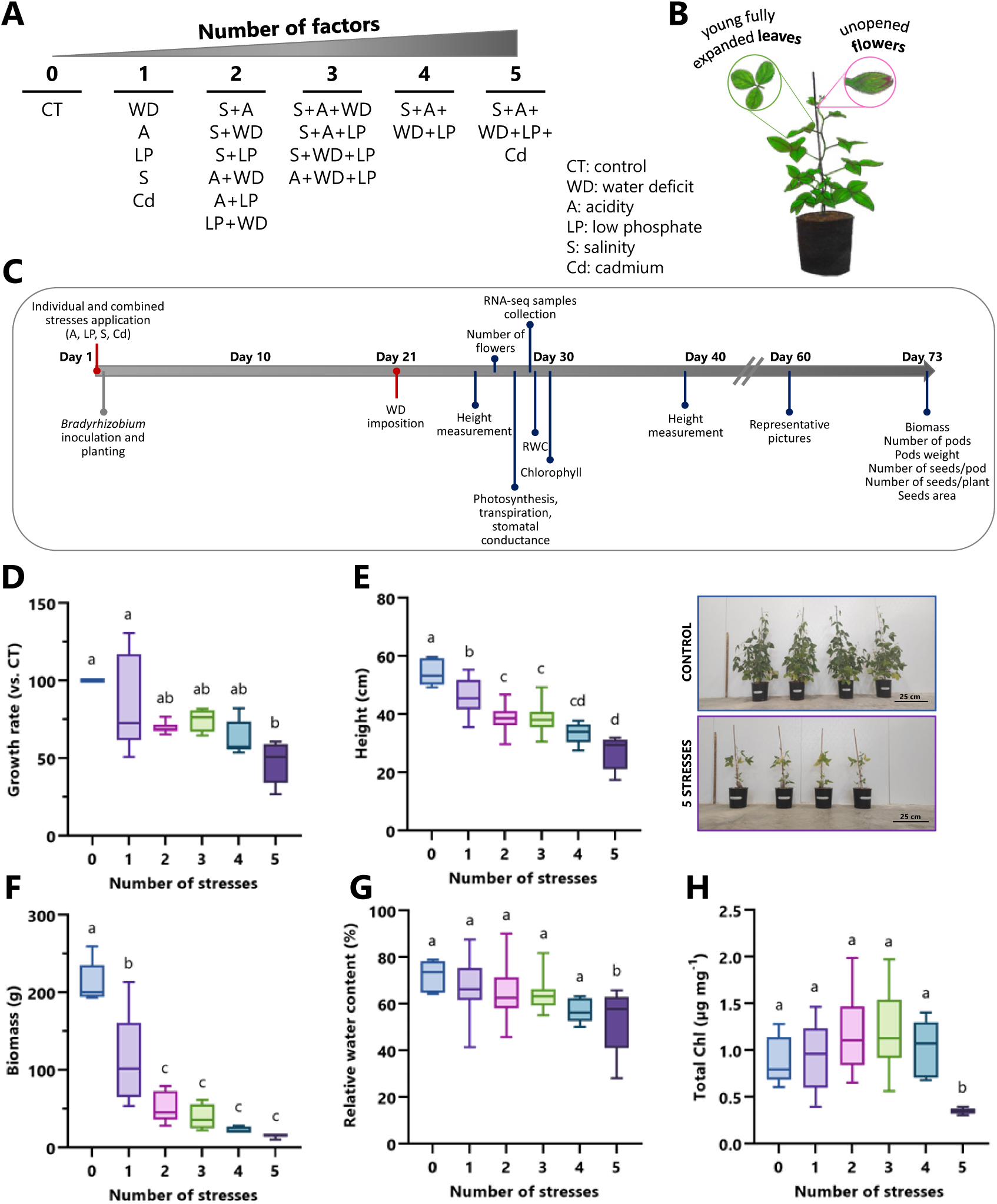
The effects of multifactorial stress combination (MFSC) on soybean growth and biomass. **A.** The different stress combinations used in the study. **B.** The different tissues sampled from soybean plants. **C.** The timeline used for the different phenotypic and molecular analyses. **D.** The effects of MFSC on soybean growth rate. **E.** Left: The effects of MFSC on plant height. Right: a representative photo of plants grown under controlled growth conditions or subjected to a MFSC of 5 different stresses. **F.-H.** The effects of MFSC on soybean biomass (F), relative water content (G), and total Chlorophyll (H), respectively. See Supplemental Figures S1, S2 for graphs including all different stress conditions. Experiments were repeated in 3 biological repeats, each with 5-7 technical repeats. Results are shown as box-and whisker plots with borders corresponding to the 25^th^ and 75^th^ percentiles of the data. Statistical analysis was performed using one-way ANOVA followed by Tukey’s *post hoc* test. Different letters denote statistical significance at *P* < 0.05. Abbreviations: A, acidity; Cd, cadmium; Chl, chlorophyll; CT, control; LP, low phosphate; S, salinity; WD, water deficit.

### Physiological responses of soybean plants to MFSC

To evaluate the impacts of MFSC on soybean physiology, we measured net photosynthesis, transpiration, and stomatal conductance of leaves from plants subjected to the same MFSCs shown in Figure 1. As shown in Figure 2A-2C, net photosynthesis, transpiration, and stomatal conductance of leaves from plants subjected to MFSC gradually declined with the increase in the number and complexity of stresses applied as part of the MFSC (see also Supplemental Figure S3). These findings demonstrate that, like the findings reported for tomato (Pascual et al., 2023), MFSC has a cumulative negative impact on soybean physiology (*i.e.,* net photosynthesis, transpiration, and stomatal conductance), potentially affecting the ability of source tissues, such as leaves, to support reproductive processes, and affecting yield.

**Figure 2.**
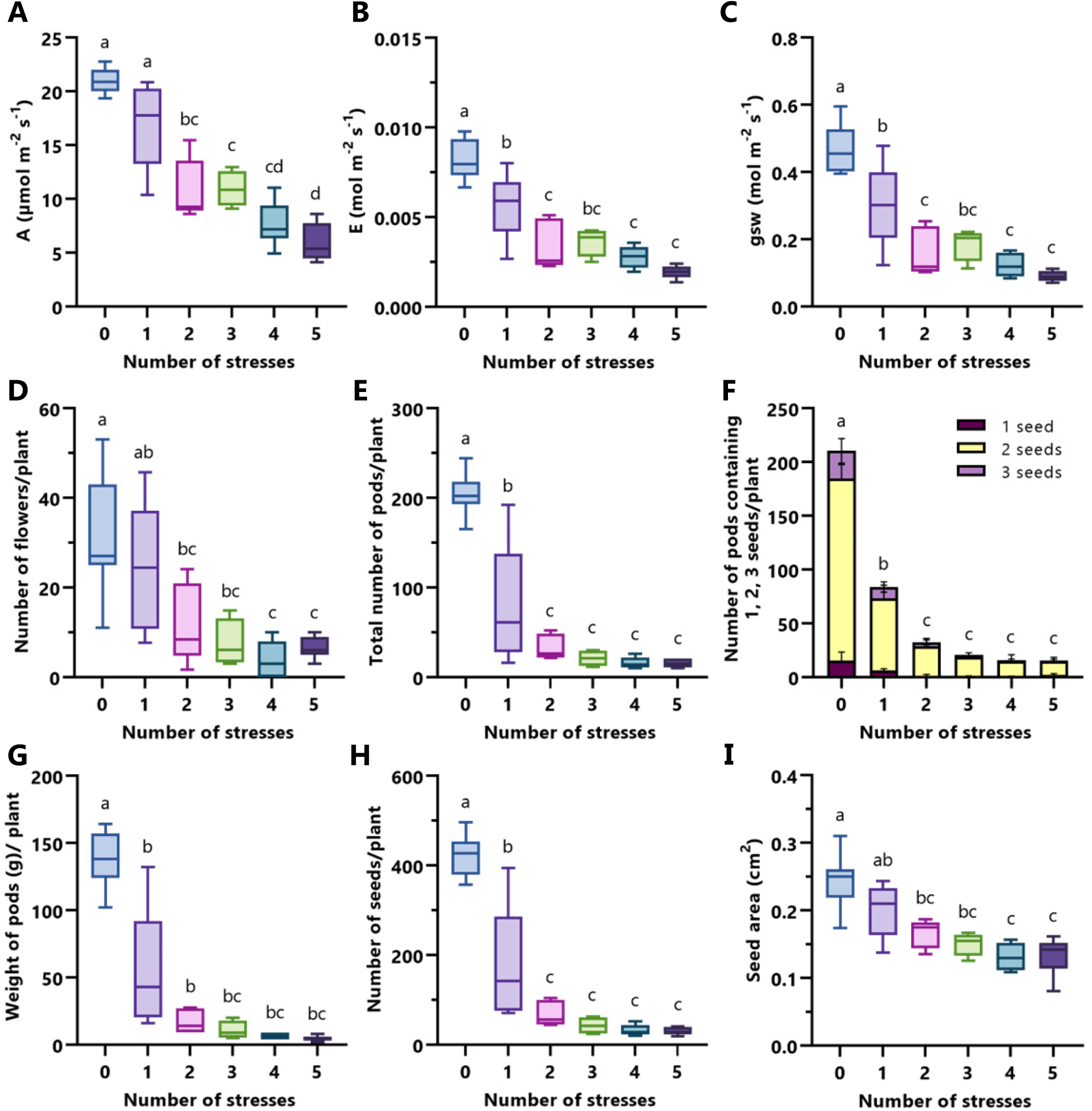
The effects of multifactorial stress combination (MFSC) on soybean physiology and reproductive parameters. **A.-C.** The effects of MFSC on soybean leaf photosynthesis (A), transpiration (B), and stomatal conductance (C), respectively. **D.-F.** The effects of MFSC on the number of flowers (D), pods (E), and pods containing 1, 2, or 3 seeds (F) of soybean plants, respectively. **G.-I.** The effects of MFSC on the weight of pods (G) and seeds per plant (H), and on seed area (I) of soybean, respectively. See Supplemental Figures S3, S4 for bar graphs including all different stress conditions. Experiments were repeated in 3 biological repeats, each with 5-7 technical repeats. Results are shown as box-and whisker plots with borders corresponding to the 25^th^ and 75^th^ percentiles of the data. Statistical analysis was performed using one-way ANOVA followed by Tukey’s *post hoc* test. Different letters denote statistical significance at *P* < 0.05. Abbreviations: A, photosynthesis; E, transpiration; gsw, stomatal conductance.

### The impact of MFSC on soybean reproductive parameters

The effects of MFSC on the yield of crop plants are largely unknown. We therefore measured different reproductive parameters of soybean plants subjected to MFSC. As shown in Figure 2D-2I, and Supplemental Figure S4, MFSC had a significant cumulative negative impact on the number of flowers and pods per plant, the number of seeds per pod, total weight of pods (including seeds) per plant, number of seeds per plants, and seed area. Taken together with the results shown in Figure 1, the impacts of MFSC on soybean plants was extensive in terms of plant growth, physiology, and reproductive success. In contrast, however, and unlike Arabidopsis (Zandalinas et al., 2021b), the MFSC conditions applied to soybean in this study did not result in plant lethality and therefore did not affect overall survival of plants (Supplemental Figure S5).

### Transcriptome of soybean flowers and leaves from plants subjected to MFSC

To begin dissecting the molecular responses of soybean to MFSC (Figure 1) and to link specific genes to the effects of MFSC on plant physiology, growth, and reproductive success (Figures 1, 2), we conducted an RNA-Seq analysis of whole flowers and leaves from soybean plants subjected to MFSC in all possible 1-, 2-, 3- and 4-combinations (as well as with the addition of cadmium as a 5^th^ stress; see below; Supplemental Data Sets 1 and 2). As shown in Figure 3A, salinity had the most robust effect on the transcriptomic response of soybean leaves, followed by water deficit, acidity, and low phosphate. Salinity and acidity, and salinity and water deficit, displayed a common response totaling over 2,400 transcripts, and all single stresses shared only 371 transcripts in common (Figure 3A; Supplemental Data Set 3). In contrast to leaves (Figure 3A; Supplemental Data Set 3), water deficit had the most robust effect on the transcriptome of whole flowers, followed by acidity, low phosphate and salinity (Figure 3B; Supplemental Data Set 4). Water deficit and acidity shared over 2,000 transcripts in common, and all single stresses shared only 271 transcripts in common. These findings suggest that the effects of each individual stress on flowers or leaves was unique, and that flowers and leaves had a very different transcriptomic response to the different stresses. Comparing the transcriptomic responses of flowers and leaves to all the different 2-stress combinations, revealed that while leaves had a relatively similar response to all 2-stress combinations (sharing over 7,200 transcripts; Figure 3A), whole flowers did not, and only 263 transcripts were common to the transcriptomic response of flowers to all 2-stress combinations (Figure 3B). A similar difference was observed between the transcriptomic response of flowers and leaves to all 3- and 4-stress combinations, with leaves showing over 9,100 transcripts in common, while flowers showing only over 1,100 (Figure 3). The findings presented in Figure 3 reveal that whole flowers had a very different transcriptomic response to MFSC compared to whole leaves and that, overall, the response of flowers appears to be more stress specific, compared to leaves.

**Figure 3.**
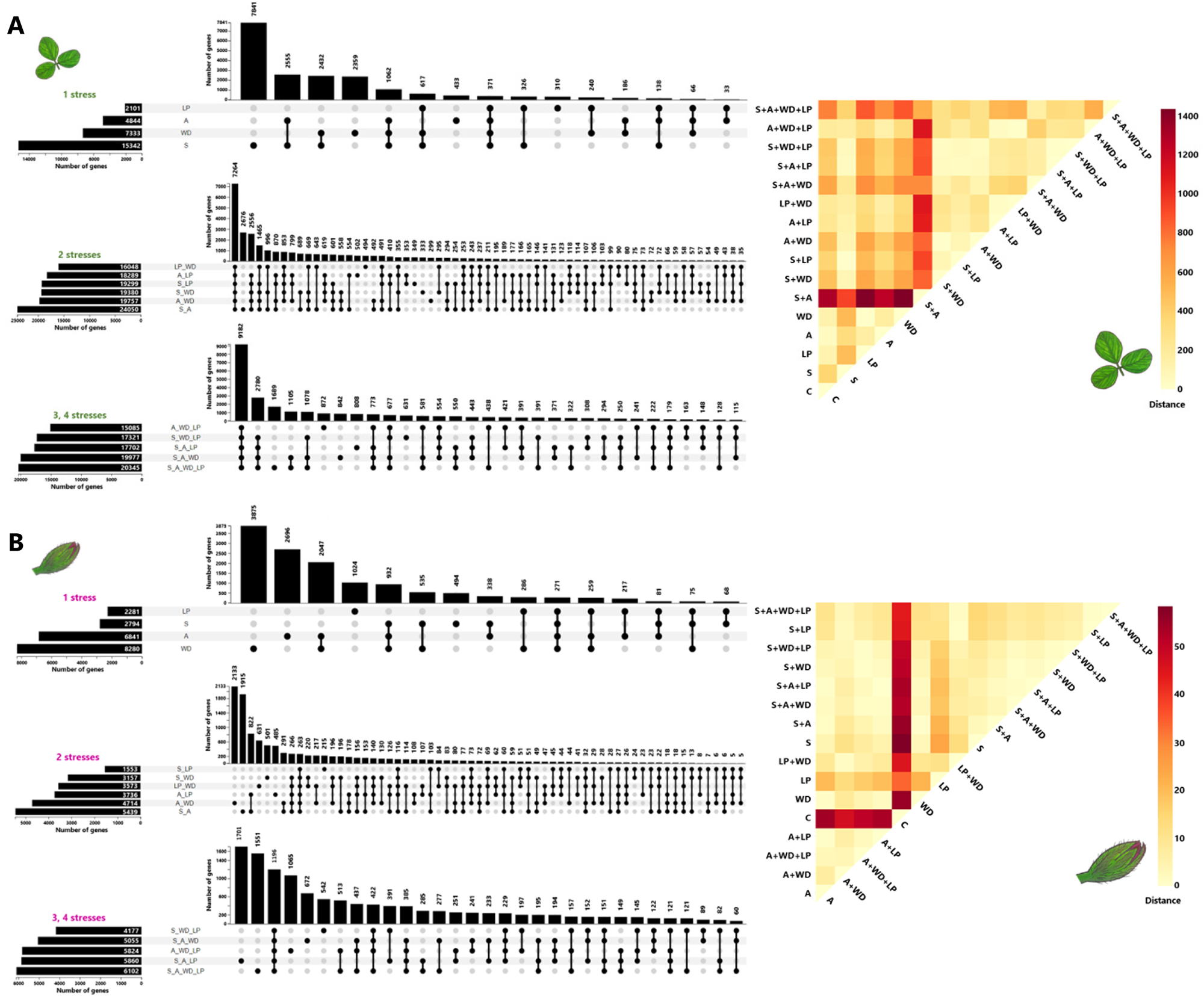
The transcriptomic response of soybean leaves and flowers to MFSC. **A.** Left: UpSet plots showing the overlap between transcripts significantly altered in soybean leaves in response to all 1-, 2-, 3-, and 4-stress combinations. Right: A distance matrix for the response of soybean leaves to MFSC of up to 4 different stresses. **B.** Same as in A, but for soybean flowers. Experiments were repeated in 3 biological repeats, each with a pool of 5-7 technical repeats. Differentially expressed transcripts were defined as those that had an adjusted *P* < 0.05 (negative binomial Wald test followed by Benjamini–Hochberg correction). A Principal Component Analysis is shown in Supplemental Figure S6. Abbreviations: A, acidity; CT, control; LP, low phosphate; S, salinity; WD, water deficit.

To better understand the relationship(s) between the transcriptomic responses of the two soybean tissues (flowers and leaves) to the different stresses/stress combinations, we generated distance matrixes for these responses. As shown in Figure 3, the combination of salinity and acidity had the most unique effect on the leaf transcriptome (Figure 3A, right panel), while the effects of each of the different stresses/stress combinations was highly different than control in flowers (Figure 3B, right panel). These findings support the results shown in the UpSet plots presented in Figure 3, left panels, demonstrating that a combination of salinity and acidity elicits the most unique leaf transcriptomic response, as well as further revealing that, compared to leaves, the response of flowers to each stress/stress combination involved in the MFSC is unique.

To directly compare the transcriptomic responses of soybean flowers and leaves to MFSC, we generated Venn diagrams comparing the transcripts common to the response of flowers and leaves to all 1-, 2-, 3-, and 4-stress combinations, and their overlap (Figure 4A, 4B, Supplemental Data Set 5). We first examined the overlap between all 1-, 2-, and 3-stresses/stress combinations of flowers and leaves. As shown in Figure 4A (see also Supplemental Data Set 5), there was very little overlap between all transcripts common to these stresses/stress combinations within each tissue and between the different tissues (the one gene in common was GLYMA_09G053700, Ankyrin repeat family protein). We next tested the overlap between all the transcripts common to all 1-, 2-, and 3-stresses/stress combinations, and the 4-stress combination in flowers and leaves. As shown in Figure 4B, a very low degree of overlap could also be found in this comparison, with only one gene common to all conditions (the same GLYMA_09G053700, Ankyrin repeat family protein). Taken together, the results presented in Figures 3 and 4 demonstrate that the response of soybean flowers and leaves to each stress/2-stress combination/MFSC is different, and that the responses of soybean leaves and flowers to these conditions are different from each other.

**Figure 4.**
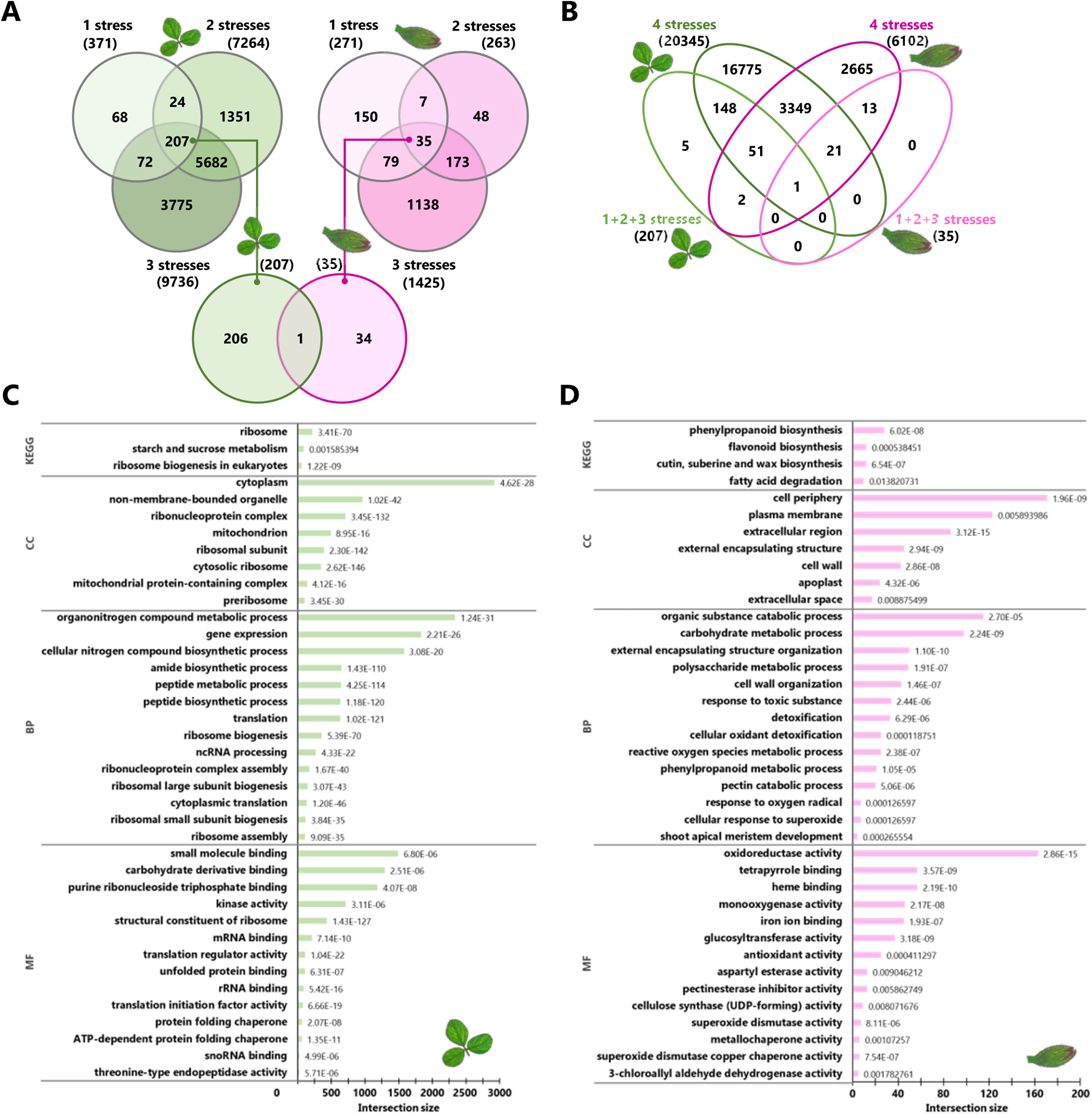
Venn diagrams and GO/KEGG analyses of soybean transcripts significantly altered during MFSC. **A.** and **B.** Venn diagrams comparing the transcripts common to the response of soybean flowers and leaves to all 1-, 2-, and 3-stress combination (A), and 1+2+3 and 4-stress combinations (B), and their overlap. **C.** KEGG and GO annotation analyses of the different transcripts identified in the soybean leaf transcriptomic response to all 3- and 4-stress combinations. **D.** Same as in C, but for soybean flowers. Complete GO/KEGG results are shown in Supplemental Data Set 6. Numbers in bars in panels C and D depict p-values. Abbreviations used: BP, biological process; CC, cellular component; GO, gene ontology; KEGG, Kyoto encyclopedia of genes and genomes; MF, molecular function.

### Enrichment term analyses of the response of soybean plants to MFSC

To further study the response of soybean leaves and flowers to MFSC, we pooled the transcripts significantly altered in their expression in all 3- and 4-stress combinations and conducted a gene ontology (GO) analysis on them. As shown in Figure 4C and Supplemental Data Set 6, among the most common terms that correlate with the transcriptomic response of soybean leaves to MFSC were terms such as ‘ribosome function’, ‘ribosome structure’, ‘translation’, and ‘RNA processing’. A similar result was found in a Kyoto Encyclopedia of Genes and Genomes (KEGG) analysis of this data set (Figure 4C). In contrast, a GO/KEGG enrichment analysis of the transcriptomic response of flowers to MFSC revealed that it was enriched in several different processes associated with ROS metabolism, metal (iron and copper) binding, oxidoreductase and glucosyltransferase activity, phenylpropanoid metabolism, apoplast and cell wall functions, and other processes that resembled the response of plants to toxic intermediates/pathogens/stress (Figure 4D; Supplemental Data Set 6). These findings shed new light on the transcriptomic response of soybean leaves to MFSC, suggesting that it primarily involves basic modifications at the level of protein translation, as opposed to the response of flowers to MFSC that appears to involve active scavenging/ detoxification of different stress intermediates.

### The impact of 5-stress MFSC on soybean

To further expand our analysis of MFSC in soybean, we added to the 4-stress MFSC a fifth stress (the heavy metal cadmium; Figure 1). As shown in Figure 5A and Supplemental Data Set 7, the transcriptomic response of flowers to cadmium was more robust (over 4,200 transcripts), compared to leaves (2,807 transcripts). In addition, while in leaves cadmium and salinity shared many transcripts in common, in flowers, cadmium and water deficit and/or acidity shared the most transcripts in common (Figure 5A).

**Figure 5.**
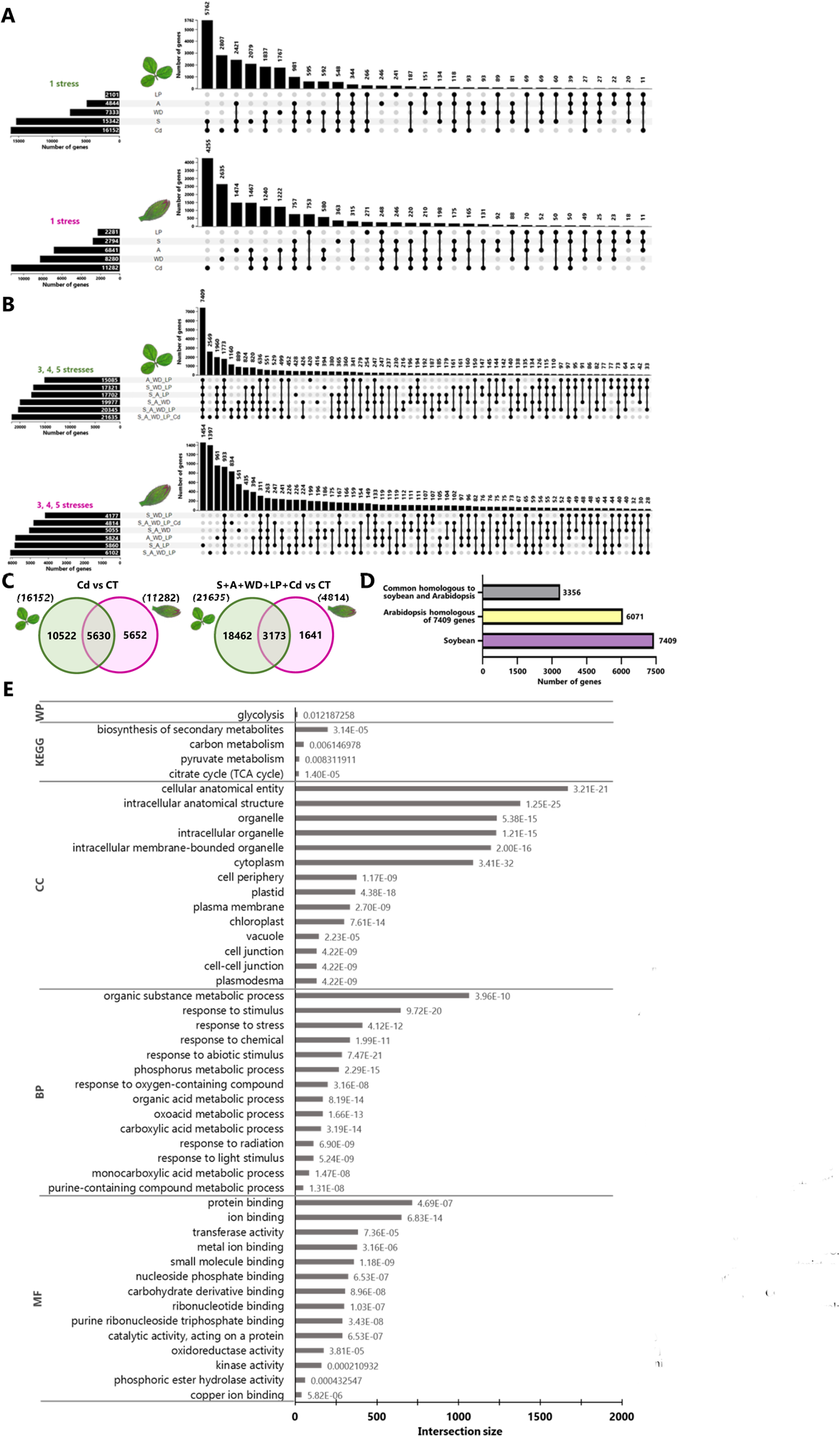
The impact of a 5-stress MFSC on the soybean transcriptome and comparison to the MFSC response of Arabidopsis. **A.** UpSet plots showing the overlap between transcripts significantly altered in soybean leaves and flowers in response to all 5 single stresses. **B.** UpSet plots showing the overlap between transcripts significantly altered in soybean leaves and flowers in response to all 3-, 4-, and 5-stress combinations. **C.** Venn diagrams comparing the response of soybean leaves and flowers to cadmium (left), or the 5-stress combination (right). **D.** Bar graph showing a comparison between all transcripts significantly altered in Arabidopsis seedlings and soybean leaves in response to MFSC. **E.** GO and KEGG annotation analysis of the genes common to the response of Arabidopsis and soybean to MFSC. Numbers in bars in panel E depict *p*-values. Abbreviations: A, acidity; BP, biological process; CC, cellular component; Cd, cadmium; CT, control; GO, gene ontology; KEGG, Kyoto encyclopedia of genes and genomes; LP, low phosphate; MF, molecular function; S, salinity; WD, water deficit; WP, wiki pathways

When comparing the transcriptomic response of soybean leaves and flowers to 3-, 4-, and 5-stress combinations (Figure 5B; Supplemental Data Set 7), it was found that many transcripts were common to the response of leaves to 3-, 4-, and 5-stresses (over 7,400 transcripts), while fewer transcripts (933) were common to the response of flowers to 3-, 4-, and 5-stresses (similar to the results shown in Figure 4 for 2- or 3- and 4-stress combinations). A Venn diagram directly comparing the 5-stress combination response of flowers and leaves revealed over 3,100 transcripts in common (Figure 5C; Supplemental Data Set 7), roughly similar to the overlap between the 4-stress combination response of flower and leaves (Figure 4). By comparison, 5,630 transcripts were common to the leaf and flower response to cadmium alone (Figure 5C; Supplemental Data Set 7). A GO/KEGG analysis with the transcripts significantly altered in their expression in response to all 3-, 4- and 5-stresses revealed similar results to the 3- and 4-stress combinations shown in Figures 4C, 4D and Supplemental Data Set 6 (Supplemental Data Set 8).

To study how conserved the transcriptomic response of soybean leaves (Figures 3-5) and Arabidopsis seedlings (Zandalinas et al., 2021b) is to MFSC, we took the list of all soybean transcripts common to 3-, 4-, and 5-stress combinations (7,409; Figure 5B) and used the OMA Browser genome pair orthology tool (Altenhoff et al., 2021) to identify their Arabidopsis homologs (we identified 6,071 Arabidopsis transcripts in this list; Supplemental Data Set 9). We then tested how many of these 6,071 homologs were found in the list of all transcripts common to the response of Arabidopsis seedlings to 3-, 4-, 5-, and 6-stress combinations (3,512 transcripts; Zandalinas et al., 2021b). This analysis revealed that 3,356 transcripts were common to the response of soybean leaves and Arabidopsis seedlings to MFSC (Figure 5D). A GO/KEGG enrichment term analysis of this conserved group of transcripts revealed that it was enriched in transcripts involved in metal and ion binding, oxidoreductase activity, and responses to stress, radiation, chemicals, and oxygen-containing compounds (Figure 5E; Supplemental Data Set 9).

The findings presented in Figure 5D-E suggest that although different MFSCs elicit relatively unique responses within each plant (Figures 3-5; Zandalinas et al., 2021b), these responses share common cellular functions/processes, primarily involved in the detoxification of different stress-related compounds and metal binding. These functions, and the transcripts/genes controlling them, could represent promising targets for biotechnological/breeding interventions in different crop plants, aimed at enhancing their tolerance to stress, stress combinations, and MFSC (Zandalinas and Mittler 2022; Rivero et al., 2022).

### Expression of stress-response transcription factors in leaves and flowers of soybean plants subjected to MFSC

To begin dissecting the transcriptional responses of soybean to MFSC, we focused on three different families of transcription factors (TFs): heat shock transcription factors (HSFs), dehydration responsive element binding (DREB) TFs, and related to AP-2 Group VII ethylene response factors (RAP) TFs (Zandalinas et al., 2020b). As shown in Figure 6A, while the steady-state level of several transcripts encoding HSFs (HSFB1, HSFA8, and HSFB2A) was enhanced by all 3-, 4-, and 5-stress combinations in leaves, the steady-state level of these TFs was not enhanced by these MFSCs in flowers. Differences in the steady-state transcript level of DREBs (and some of their associated transcripts), and RAPs, were also identified between leaves and flowers of soybean plants subjected to MFSC (Figure 6A). Overall, it appears that the expression level of many of these stress-associated TFs (HSFs, DREBs, and RAPs) was altered more vigorously in leaves compared to flowers (Figure 6A), perhaps indicating that flowers experience less stress, or use other stress-response TFs, compared to leaves, under conditions of MFSC. In this respect, it should be mentioned that our comparative analysis of flowers and leaves from soybean plants subjected to drought, heat stress, and their combination (Sinha et al., 2022b, 2023a, 2023b) reveled a somewhat similar result, suggesting that flowers (and pods; Sinha et al., 2023a, 2023b) are better protected from stress, compared to leaves.

**Figure 6.**
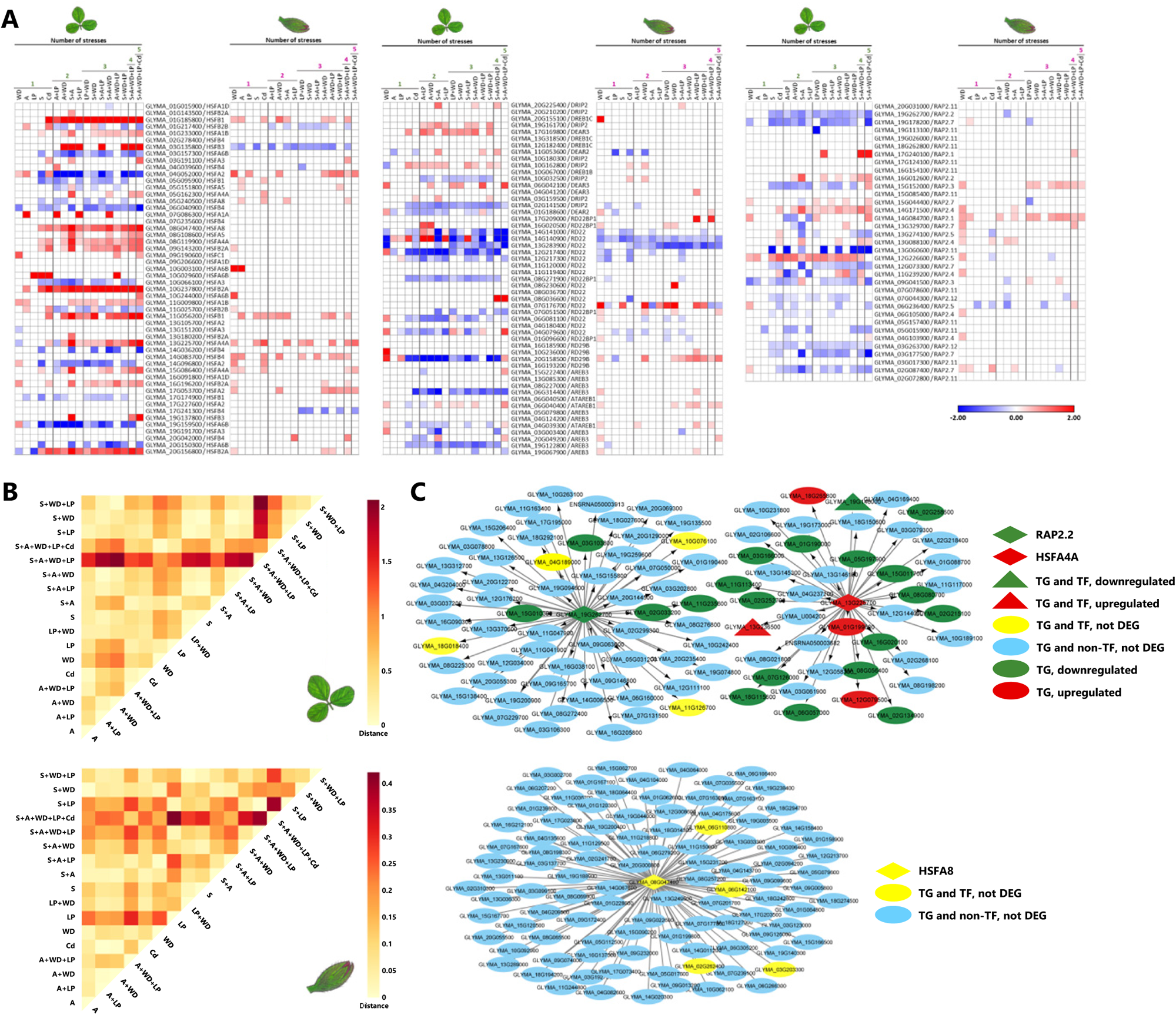
Expression of selected transcription factors in leaves and flowers of soybean plants subjected to MFSC. **A.** Heatmaps for the expression of soybean HSF, RAP, and DREB TFs in flowers and leaves during MFSC. **B.** Distance matrixes for the top 100 transcripts identified by a GENIE3 network analysis of RAP2.2 (GLYMA_19G262700) during the response of soybean leaves (top) or flowers (bottom) to MFSC. **C.** Representative GENIE3 network analysis results for some of the selected TFs shown in A in soybean leaves (top) or flowers (bottom) in response to the 5-MFSC condition. Upregulated transcripts are in red, downregulated transcripts are in green (log2 fold change is more or less than 0, respectively), and non-differentially expressed transcripts are in yellow. TF-encoding transcripts are represented by triangles, differentially expressed transcripts were represented by diamonds, ellipse shape transcripts represent target genes, and triangles represent target genes that are TFs. Abbreviations: A, acidity; Cd, cadmium; CT, control; DEG, differentially expressed gene; DREB, dehydration responsive element binding; HSF, heat shock factor; LP, low phosphate; RAP, related to AP-2 Group VII ethylene response factors; S, salinity; TF, transcription factor; TG, target gene; WD, water deficit.

To further dissect the transcriptional networks under the control of HSFs and RAPs, we focused on four HSFs (GLYMA_08G047400/HSFA8, GLYMA_08G119900/HSFA4A, GLYMA_10G066100/HSFA3, and GLYMA_13G225700/HSFA4A) and three RAPs (GLYMA_14G171500/RAP2.4, GLYMA_12G226600/RAP2.5, and GLYMA_19G262700/ RAP2.2), and conducted a GENIE 3 analysis (Sinha et al., 2023b) for these TFs in leaves and flowers in response to all stresses/stress combinations (Supplemental Data Sets 10 and 11). We further generated a distance matrix that compared the expression level differences of the top 100 genes regulated by RAP2.2 under all stress conditions. As shown in Figure 6B, the transcriptional responses controlled by RAP2.2 under the different stress conditions were mostly different from each other. Similar results were found for GLYMA_08G047400/HSFA8, GLYMA_10G066100/HSFA3, GLYMA_13G225700/ HSFA4A, GLYMA_08G119900/HSFA4A, GLYMA_14G171500/RAP2.4 and GLYMA_12G226600/RAP2.5, (Supplemental Data Sets 12 and 13; representative regulatory maps are shown in Figure 6C). These results demonstrate that the molecular responses of soybean flowers and leaves to the different stress conditions involved in the MFSC is mostly different from each other, even if they are associated with/regulated by the same TF. In addition, they show that, at least in leaves, the 5-stress combination is the most different when it comes to the networks associated with RAP2.2 (Figure 6B), and that the extent of the transcriptional responses regulated by this TF might be more vigorous in flowers, compared to leaves (*e.g.,* Figure 6C).

### Integrative MultiOmics Pathway Resolution (IMPRes) analysis of MFSC in soybean

To identify unique pathways associated with the response of soybean flowers to MFSC, we further applied IMPRes, a stepwise active pathway detection method that uses a dynamic programming approach, together with existing pathway interaction sources (*e.g.,* KEGG, protein-protein interaction, and other databases), to reconstruct putative networks, using transcriptomics datasets (Jiang et al., 2020a, 202b). For this analysis we used a set of seed transcripts associated with the MFSC response of flowers to 3-, 4- and 5-stresses (Supplemental Data Set 14). As shown in Figure 7A, this analysis identified ascorbate and aldarate metabolism, as well as tryptophan and tyrosine metabolism, as associated with the MFSC responses of soybean flowers. The identification of ascorbate, an important antioxidant (Smirnoff, 2018; Mittler et al., 2022), is in agreement with the GO/KEGG analysis of transcripts significantly altered in their expression in flowers during MFSC (Figure 4D; Supplemental Data Sets 6 and 8), as well as our previous results in Arabidopsis (Zandalinas et al., 2021b), showing that tolerance to MFSC is significantly compromised in Arabidopsis mutants deficient in ROS scavenging (Ascorbate peroxidase 1; *apx1*) and ROS signaling (Respiratory burst oxidase homolog D; *rbohD*), further supporting the IMPRes analysis shown in Figure 7A.

**Figure 7.**
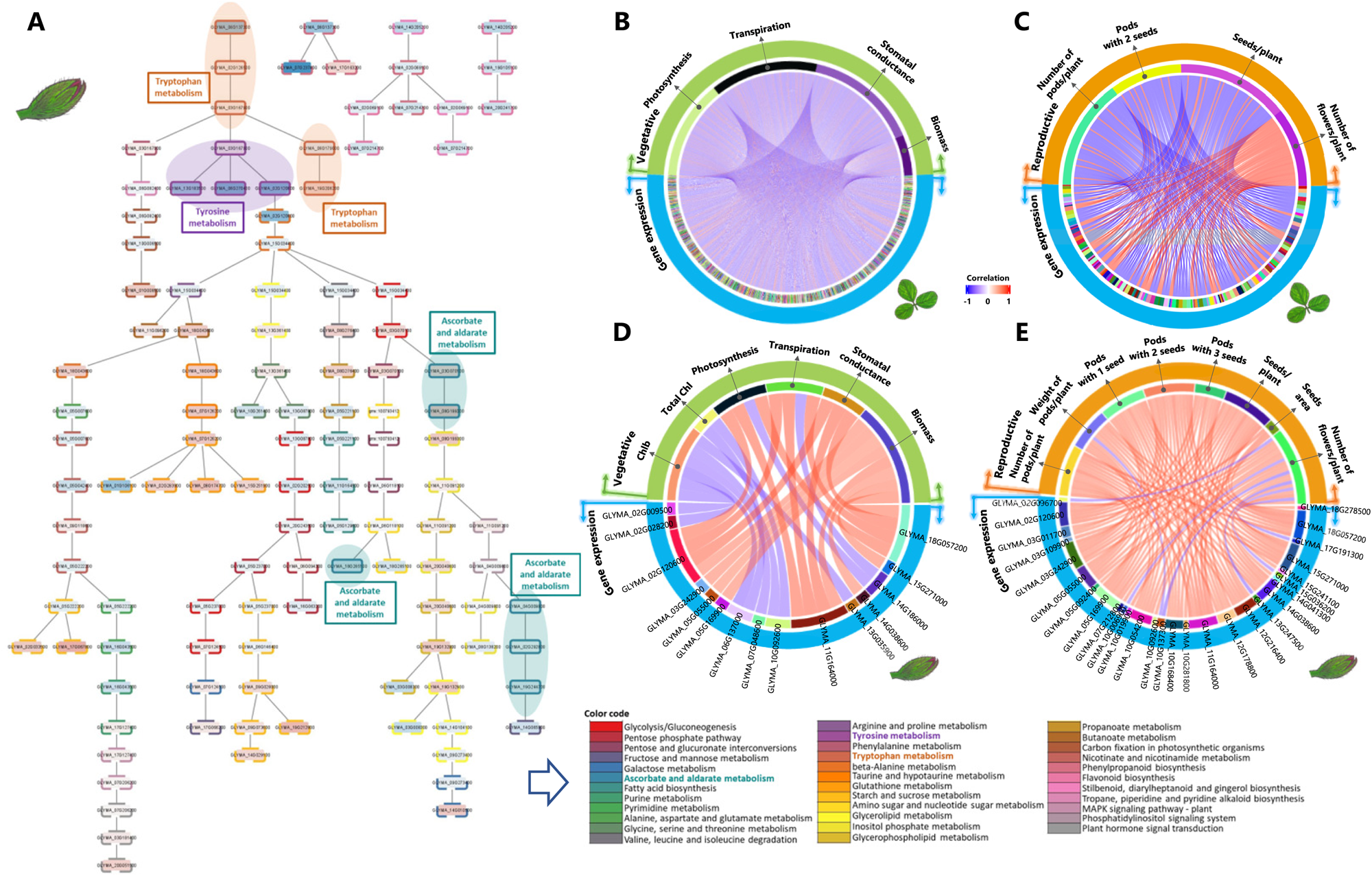
IMPRes and mixOmics analyses of MFSC in soybean. **A.** A representative pruned network diagram for the IMPRes *in silico* pathway analysis of the response of soybean flowers to MFSC. Seed transcripts used for the analysis are shown in Supplemental Data Set 14. Transcripts are represented by nodes and their expression levels are indicated by the colors red and blue, which correspond to upregulation and downregulation, respectively. Each pathway in the diagram is assigned a distinct color. The significance of each transcript within a pathway is depicted by the border pattern surrounding the node. Transcripts that are deemed significant (*p* < 0.05) are depicted with a solid border, non-significant genes have a dashed border. **B.-E.** MixOmics analysis linking specific transcripts expressed in leaves (B, C) or flowers (D, E), with specific vegetative (B, D) or reproductive (C, E) phenotypes. The transcriptomics data used is the combination of differentially expressed genes (DEGs) identified with *q*-value less than 0.01 and log2 fold change of 2 or more for all stress comparisons. This cutoff extracted 5913 DEGs for leaf group and 464 DEGs for flower group. The reproductive data used include number of pods, weight of pods, number of pods with 1, 2, or 3 seeds, weight of seeds, and number of flowers, per plant, as well as seed area; the vegetative data used include growth rate, relative water content, biomass, Chl a, Chl b, and total Chl; and the physiological data used include photosynthesis, transpiration, and stomatal conductance. Abbreviation used: Chl, chlorophyll; DEG, differentially expressed gene; IMPRes, Integrative MultiOmics Pathway Resolution.

### Integrating reproductive success, physiology, and plant growth with transcriptomic data using mixOmics

To begin identifying links between the transcriptomic responses of soybean flowers and leaves (Figures 3-7A), and vegetative (*e.g.,* plant growth, biomass, and physiology; Figures 1D-1H, 2A-2C; Supplemental Figures S1-S3), and reproductive (*e.g.,* number of flowers, pods, seeds per pod; Figure 2D-2I; Supplemental Figure S4) phenotypes of soybean, during MFSC, we utilized the multivariate tool mixOmics (Rohart et al., 2017). As inputs we used all transcripts altered in response to all 1-, 2- and all MFSC treatments (Figures 3-7A; Supplemental Data Sets 1 and 2), all physiological and growth parameters (Figures 1D-1H, 2A-2C, Supplemental Figures S1-S3), and all reproductive success parameters (Figures 2D-2I, Supplemental Figure S4). As shown in Figure 7B-7E and in Supplemental Data Set 15, this analysis yielded putative interactions between specific genes expressed in leaves (Figure 7B, 7C) or flowers (Figure 7D, 7E), during MFSC, and specific vegetative (Figure 7B, 7D) or reproductive (Figure 7C, 7E) phenotypes. Among the flower-expressed transcripts associated with reproductive success parameters were ROS-response transcripts such as copper/zinc superoxide dismutase (Cu/Zn-SOD) 1 and 2, Cu/Zn-SOD copper chaperone, copper amine oxidase, and glutathione-S-transferases, as well as GDSL-like Lipase, laccase, beta glucosidase, and xyloglucan endotransglucosylase (Supplemental Data Set 15). Among the leaf-expressed transcripts associated with yield parameters were different TFs (*e.g.,* WRKY, NAC, KNOTTED, and bHLH), receptors, channels/transporters, disease resistance and programmed cell death related proteins, kinases, and proteins involved in stomata and plasmodesmata development/functions (Supplemental Data Set 15). Included with the leaf-expressed transcripts associated with vegetative phenotypes were different kinases and phosphatases, receptors, peroxidases, TFs such as WRKY, and many different proteins involved in multiple functions (Supplemental Data Set 15). The leaf and flower transcripts associated with reproductive success parameters, and the leaf transcripts associated with vegetative growth (Supplemental Data Set 15), represent promising leads for biotech interventions attempting to enhance the tolerance of soybean plants to MFSC (Zandalinas and Mittler 2022; Rivero et al., 2022).

## DISCUSSION

Global warming, and the different disruptions it causes to our climate, coupled with the accumulation of various man-made pollutants, the degradation of soils and microbiome diversity in many parts of our globe, and the climate-driven changes in pathogen/weeds/invasive species activities, are likely to subject crop plants to different combinations of MFSC, in the coming years (Masson-Delmotte et al., 2021; Rillig et al., 2019; Zandalinas et al., 2021a; Zandalinas and Mittler, 2022). Recent studies conducted in Arabidopsis, tomato, rice, maize, and now soybean, demonstrate that with the increasing number and complexity of MFSCs affecting crops, the growth, survival, and/or yield of crops declines (Figures 1, 2; Zandalinas et al., 2021b; Sinha et al., 2022a; Pascual et al., 2023), highlighting the importance of studying MFSC with the goal of increasing the tolerance of plants to this impending threat (Zandalinas et al., 2021a; Zandalinas and Mittler, 2022; Rivero et al., 2022). While, to the best of our knowledge, no comprehensive study has been conducted so far on field-grown crops subjected to MFSC, several studies have shown that MFSC has a severe negative impact on soils, soil microbiomes, forests, lagoons, and other ecosystems (Adams et al., 2019; Popkin, 2021; Rillig et al., 2019, 2023; Yang et al., 2022), further supporting the need to study MFSC in crops (Zandalinas et al., 2021a, 2021b; Mittler and Zandalinas, 2022). In the current study we report that MFSC has a significant negative impact on the growth, physiology, and reproductive success of the globally important crop plant, soybean (Figures 1, 2). We further provide unique and integrative transcriptomic and phenotypic datasets for the response of soybean flowers and leaves to MFSC (Figures 3-7; Supplemental Figures S1-S4; Supplemental Data Sets 1-15).

Comparing the transcriptomic responses of flowers and leaves to MFSC revealed significant differences between these two plant tissues. While flowers appear to alter the expression of many cellular detoxification transcripts in response to MFSC (Figure 4D; Supplemental Data Set 2), leaves had a different transcriptomic response to MFSC that included alterations in ribosome structure/function and changes in translation (Figure 4C; Supplemental Data Set 1). This finding could suggest that the stress level flowers and leaves experience during MFSC is different, or that the response of flowers to MFSC is distinct from that of leaves. In two recent studies of the physiological and molecular responses of soybean flowers and leaves to a combination of drought and heat stress (Sinha et al., 2022b, 2023b), it was found that flowers (as well as pods; Sinha et al., 2023a) were more protected from the effects of the stress/stress combination compared to leaves. In addition, it was found that active transpiration is maintained in flowers and pods during the stress combination enabling the cooling of these tissues, while transpiration was suppressed in leaves (termed ‘differential transpiration’; Sinha et al., 2022b, 2023a), subjecting leaves to a higher internal temperature. Taken together, these findings suggest that compared to leaves, flowers are more protected during MFSC, perhaps because they represent an important sink tissue required for the reproduction of an annual plant like soybean (Sinha et al., 2022b). In future studies it would be important to measure the stomatal conductance, index, transpiration, and other physiological, biochemical, and molecular processes occurring in flowers and pods (Sinha et al., 2022b, 2023a), as well as determine the transcriptomic responses of different flower and pod tissues (Sinha et al., 2023b), and single cells within these tissues (Cervantes-Pérez et al., 2022), to MFSC. These studies could provide more promising leads for biotechnological interventions in soybean and other crops attempting to make them more tolerant to MFSC (Zandalinas and Mittler 2022; Rivero et al., 2022). When comparing the transcriptomic responses of flowers and leaves to the individual abiotic stresses, it is also interesting to note that soybean flowers and leaves were affected differently by water deficit, acidity, and salinity (Figure 4). Whereas flowers had a more robust transcriptomic response to water deficit or acidity, leaves had a more robust transcriptomics response to salinity. These differences support the notion that flowers are better protected from certain stresses (*e.g.,* salinity), but are more prone to other stresses, such as water deficit, perhaps due to the enhanced transpiration occurring in flowers during stress (Sinha et al., 2022b).

To link specific genes expressed in flowers and leaves to the effects of MFSC on different reproductive success, growth, and physiological parameters, we used the mixOmics platform (Figure 7B-E). Our analysis revealed that the expression of many ROS-response transcripts (*e.g.,* Cu/Zn-SODs and their copper chaperons) in flowers is associated with the effects of stress/MFSC on reproductive success (Supplemental Data Set 15). In contrast, the expression of many TFs and other stress- and development-associated genes in leaves was associated with the effects of MFSC on reproductive success. The impact of manipulating the expression of selected groups of these genes, in flowers and/or leaves of soybean, on yield of plants subjected to MFSC should be addressed in future studies. These studies could use CRISPR or stress/MFSC-specific expression strategies and would require lab, greenhouse, and field studies (Mittler and Blumwald, 2010). Of particular interest is the role of copper chaperones and Cu/Zn SODs in flowers (Supplemental Data Set 15). Transgenic crops ectopically expressing SODs had limited success in mitigating the effects of stress on yield of field-grown plants (Kerchev and Van Breusegem, 2022). However, in contrast to previous studies that used ectopic expression of SODs, the effects of stress-induced and/or flower-specific expression of these important superoxide scavengers on abiotic stress tolerance has not been evaluated thoroughly.

The availability of transcriptomic datasets for MFSC from two different plant species (soybean and Arabidopsis; Figure 3-7; Supplemental Data Set 9; Zandalinas et al., 2021b) enabled us to compare the response of these two species to MFSC (Figure 5C-5E). Before addressing differences between the transcriptomic responses of Arabidopsis seedlings and soybean leaves to MFSC it is important to note that while MFSC caused a significant decrease in Arabidopsis seedling survival (Zandalinas et al., 2021b), our analysis of soybean plants subjected to MFSC did not reveal a similar effect (Supplemental Figure S5). Of biotechnological and breeding interest are the transcripts common to the response of soybean and Arabidopsis to conditions of MFSC. These include responses associated with almost all subcellular compartments, as well as glycolysis, the TCA cycle, and different metal ion binding and ROS detoxification pathways (Figure 5E; Supplemental Data Set 9). In this respect it should be mentioned that seedlings of Arabidopsis mutants deficient in ROS scavenging (*apx1*), ROS signaling (*rbohD*), or the regulation of iron/iron-sulfur cluster metabolism (Arabidopsis NEET protein; *AtNEET RNAi*), were found to have reduced tolerance to MFSC in Arabidopsis (Zandalinas et al., 2021b). In addition, our IMPRes pathway analysis identified ascorbic acid (ascorbate) as playing a putative role in the MFSC response of soybean flowers (Figure 7A; Supplemental Data Set 14), and our proteomics analysis of rice seedlings subjected to MFSC revealed increased abundance of ascorbate peroxidases, catalases, SODs, and other ROS-scavenging proteins during MFSC (Sinha et al., 2022a). Taken together, it appears that ROS detoxification and/or signaling could play a key role in the response of soybean and Arabidopsis to MFSC. In future studies it would be interesting to test the tolerance to MFSC of genetically altered soybean plants with enhanced resistance to ROS stress, and/or with an enhanced ability to sequester metal ions and other reactive intermediates/metabolites that can cause the accumulation of ROS (Mittler et al., 2022). In addition, the role of balancing iron and iron-sulfur metabolism should be addressed, as iron, ascorbic acid and ROS play an important, yet dangerous, interplay in plant metabolism and stress responses (Smirnoff, 2018; Mittler et al., 2022).

In summary, our results provide unique, integrative, and large-scale biological resource datasets for future studies focusing on the impacts of MFSC on crop growth, physiology, molecular responses, and yield. Considering the overall cumulative, and mostly unabated, negative impacts of mankind on our environment (Masson-Delmotte et al., 2021), and their potential to subject major crops, trees, and other plants to MFSC (Rillig et al., 2019, 2023; Mittler and Zandalinas, 2022; Zandalinas et al., 2021a), further research is needed into this important subject. This research should include small- and large-scale field studies of the effects of MFSC on crop growth, yield, and microbiomes/soils, as well as conducting crop genetic diversity studies under conditions of MFSC, and engineering/breeding attempts to make crops more tolerant to stress combinations/ MFSC (Zandalinas and Mittler, 2022; Rivero et al., 2022). These studies are critical for mitigating the effects of current and future environmental challenges on crop yield and food security worldwide (*e.g.,* Lobell et al., 2011; Challinor et al., 2014; Mittler and Blumwald, 2010; Bailey-Serres et al., 2019).

## METHODS

All experiments were conducted in a climate-controlled greenhouse that is part of the Plant Growth Facility of the University of Missouri (https://plantgrowthfacilities.missouri.edu/). Soybean (*Glycine max*, cv *Magellan*) seeds were coated with *Bradyrhizobium japonicum* inoculum (N-DURE, Verdesian Life Sciences, NC, USA) and sown at 4 cm depth in plastic nursery pots (7.57 L, Hummert International, Inc) filled with 1.5 kg of Promix BX growing medium (Premier Tech Horticulture; PA, USA) and perlite (Miracle-Gro® Perlite, Miracle-Gro, Marysville, OH, USA) mixed at a ratio of 10:1 and soaked with 3 L of nutrient solution. The five stress conditions were acidity (pH 4; A), water deficit (30% of the water available for transpiration; WD; Sinha et al., 2022b), salinity (15 mM NaCl; S), phosphate deficiency (phosphate concentration was reduced by 90%; LP), and cadmium (300 µM CdCl_2_; Cd). To study MFSC, WD, A, LP and S stresses were conducted in all possible combinations (Figure 1A). To generate a MFSC of five factors, Cd was added to the 4-stress combination. Except for the water deficit treatment, all stresses alone or in combination were added to the nutrient solution at the start of the experiment (Figure 1A, 1C). A total of 126 pots containing one plant each were divided into 18 experimental groups with 7 replicates per treatment as described in Figure 1A. The composition of the control nutrient solution was: 149.2 mg L^-1^ KCl, 221.98 mg L^-1^ CaCl_2_, 60.19 mg L^-1^ MgSO_4_, 18.26 mg L^-1^ K_2_HPO_4_, 43.55 mg L^-1^ KH_2_PO_4,_ 9.18 mg L^-1^ FeSeq.330, 172.5 mg L^-1^ ZnSO_4_.7H_2_O, 24.2 mg L^-1^ NaMoO_4_.2H_2_O, 26.15 mg L^-1^ NiCl_2_.6H_2_O, 2.38 mg L^-1^ CoCl_2_, 37.5 mg L^-1^ CuSO_4_.5H_2_O, 142.21 mg L^-1^ H_3_BO_3_, 1.81 mg L^-1^ MnCl_2_.4H_2_O. For inducing low phosphate conditions, the nutrient solution was adjusted to 158.9 mg L^-1^ KCl, 6.1 mg L^-1^ K_2_HPO_4_, 0.68 mg L^-1^ KH_2_PO_4_, 24.4 mg L^-1^ K_2_SO_4_. For the salinity treatment, the nutrient solution was adjusted to 294 mg L^-1^ CaCl, 123.25 mg L^-1^ MgSO_4_, 876.6 mg L^-1^ NaCl. For adding the cadmium stress, the nutrient solution was adjusted to 158.9 mg L^-1^ KCl, 6.1 mg L^-1^ K_2_HPO_4_, 0.68 mg L^-1^ KH_2_PO_4_, 78.4 mg L^-1^ K_2_SO_4_, 11.42 mg L^-1^ CdCl_2_. The stock solution of each nutrient solution was prepared separately, and appropriate volumes were mixed to make up nutrient solution for all the individual and combined treatments. For plants subjected to acidity treatment, the pH of the standard nutrient solution was adjusted to 4 using H_2_SO_4_.

All plants were grown under short-day conditions (1000 µmol photons m^-2^ s^-1^, 14-h light/10-h dark) at 28/24 °C day/night temperature. Water deficit treatment alone or in combination with other stresses was imposed three weeks after planting (Figure 1C). Thereafter and throughout the experiment the plants under water deficit treatment were irrigated with only 30% of the water available for transpiration (determined by weighing pots daily as described in Sinha et al., 2022b), while plants not subjected to water deficit stress were irrigated with 100% of the water available for transpiration. Once a week, control, or low phosphate nutrient solution alone or in combination with other stresses was applied as fertilizer according to each treatment. Salinity treatment (15 mM) alone or in combination with other stresses was added just once at the beginning of the experiment as explained before. Cadmium (300 µM) as single stress or as the five-stress condition was included in the nutrient solution every two weeks. A timeline of data collection and sampling performed in this work is presented in Figure 1C. All data was collected from at least 5-7 independent replicates (depending on data type), between 9:30 AM-12 PM.

### Growth parameters

Plant height was manually measured from 7 plants/treatment at two different time points (26 and 38 days after planting; Figure 1C) and the growth rate was calculated as the difference between these two values expressed as percentage compared to control plants. Representative pictures of the plants were taken after 60 days.

### Biomass Determination

Biomass was measured at the end of the experiment (73-day-old plants) from 5 individual plants. First, the pods of each plant were collected to measure reproductive-related parameters and then the aerial part including stem and leaves was harvested. The total biomass content per plant was determined from the sum of stem and leaf tissue and expressed as grams per plant.

### Relative water content estimation

Relative water content (RWC) was measured in leaves (second fully developed leaves) from 5 plants/treatment 10 days following the start of the water deficit stress application. Fresh weight (FW) of leaf samples was measured immediately after sample collection and leaves were hydrated by floating on distilled water. Turgid weight (TW) was recorded once leaf acquired full turgidity after 5 days at 4°C. Samples were then oven dried at 60°C for 6 days, and the dried weigh (DW) was measured. RWC was calculated using the formula: RWC (%) = (FW−DW)/(TW−DW) ∗100.

### Chlorophyll content

The second fully developed leaves from 5 different plants were collected for the chlorophyll content analysis. Leaf discs (diameter 1.5 cm) obtained from the center of the leaves, avoiding the major leaf veins, were weighted to determine the fresh weight. Samples were incubated in 5 ml of *N, N*-dimethylformamide (DMF) at 4°C in the dark for 5 days. Then, the absorbance of 1 ml of the DMF extraction was read in a spectrophotometer at 603, 647 and 664 nm, using 1 ml of clean DMF as blank. Chlorophyll content was calculated as described by Moran (1982) and expressed as µg mg^-1^.

### Physiological measurements

Photosynthesis (A), transpiration (E) and stomatal conductance (gsw) parameters were measured using the second fully developed leaves of plants at two independent times (Figure 1C) using a Li-Cor Portable Photosynthesis System (LI-6800; Li-Cor, Lincoln, NE, USA) following the manufacturer’s recommendations (Sinha et al., 2022b). The results presented are the average from 5 different plants and two independent measurements.

### Reproductive parameters

Number of flowers per plant, pods per plant, seeds per pod, pods weight, total seeds per plant, and seed area were recorded for all the individual and stress combinations as described in Sinha et al., 2022b. The number of flowers per plant was manually counted 7 days after the start of the flowering phase and during 5 consecutive days, while the rest of the reproductive success parameters were measured at the end of the experiment at day 73 (Figure 1C). For each plant and treatment, pods were collected, counted, and weighted. The number of seeds per pod and the number of pods per plant were determined and then the seeds were harvested from the pods. Seed area was measured from pictures using ImageJ software. Seed pictures were taken from pods harvested from 7 different plants per treatment, and at least 30 seeds per plant were analyzed for size as described above.

### Tissue collection and RNA isolation

Sampling of flowers and leaves for the transcriptomic analyses presented in this study was conducted 10 days following the start of the water deficit treatment using new organs that developed under the stress conditions (Figure 1B-C). Samples were collected between 10 AM-12 PM. Flowers at stages II and III (unopen flowers undergoing self-pollination) were selected as described in Sinha et al., (2022b, 2023b). For each biological repeat, leaf or flowers were pooled together from 2-3 different plants (in 3 technical repeats) and flash frozen in liquid nitrogen. RNA from flower samples was isolated using the RNeasy plant mini kit (Qiagen, Germantown, MD, USA; Sinha et al., 2022b), while RNA from leaves was isolated using the RNeasy PowerPlant kit (Qiagen, Germantown, MD, USA; Sinha et al., 2022b).

### RNA sequencing and data analysis

RNA libraries for sequencing were prepared using standard Illumina protocols and RNA sequencing was performed using NovaSeq 6000 PE150 by Novogene co. Ltd (https://en.novogene.com/; Sacramento, CA, USA). Read quality control was performed using Trim Galore v0.6.4 (https://www.bioinformatics.babraham.ac.uk/projects/trim_galore/) & FastQC v0.11.9 (https://www.bioinformatics.babraham.ac.uk/projects/fastqc/). The RNA-seq reads were aligned to the reference genome for Soybean - Glycine max v2.1 (downloaded from ftp://ftp.ensemblgenomes.org/pub/plants/release-51/fasta/glycine_max/dna/), using Hisat2 short read aligner (Kim et al., 2019). Intermediate file processing of sam to sorted bam conversion was carried out using samtools v1.9 (Danecek et al., 2021). Transcript levels (expressed as FPKM values) were calculated using the Cufflinks tool from the Tuxedo suite (Trapnell et al., 2012), guided by genome annotation files downloaded from the same source. Differential gene expression analysis was performed using Cuffdiff tool (Trapnell et al., 2013), from the same Tuxedo suite. Differentially expressed transcripts were defined as those that had adjusted *P* < 0.05 (negative binomial Wald test followed by Benjamini–Hochberg correction). A Principal Component Analysis (PCA) was performed for leaf and flower samples from this study and the distance between the different replicates is shown in Supplemental Figure S6. Functional annotation and quantification of overrepresented gene ontology (GO) terms (*P* < 0.05) and KEGG enrichment were conducted using g:Profiler (Raudvere et al., 2019). Venn diagrams were created in VENNY 2.1 (BioinfoGP,CNB-CSIC) and Upset Plots were generated in upsetr (gehlenborglab.shinyapps.io; Conway et al., 2017). Morpheus software (https://software.broadinstitute.org/morpheus) was used for the heatmaps generation.

### Gene regulatory network analysis

The R package GENIE3 (1.18.0; Huynh-Thu et al., 2010), which infers a gene regulatory network (in the form of a weighted adjacency matrix) using gene expression data, was used to identify the targets of the selected transcription factors (TFs, GLYMA_08G047400/HSFA8, GLYMA_08G119900/HSFA4A, GLYMA_10G066100/HSFA3, GLYMA_13G225700/HSFA4A, GLYMA_14G171500/RAP2.4, GLYMA_12G226600/RAP2.5, and GLYMA_19G262700/RAP2.2) in leaf and flower samples using the Random Forest tree calculation method. The higher the weights, the more likely are the regulatory connections between the TFs and their targets. A weight cutoff of 0.42 and 0.49 was used for leaf and flower under S+A+WD+LP+Cd conditions, respectively, to get the top regulatory connections. This cutoff weighted adjacency matrix was later used to generate a gene regulatory network using the Cytoscape tool (3.9.1; Otasek et al., 2019) for the different TFs.

### Distance matrix

Distance matrix graphs were created using the Plotly graph objects Python package (Plotly Technologies Inc. 2021), with the distances calculated using the Euclidean distance metric (Deza and Deza, 2009). Specifically, we focused on the top 100 Differentially Expressed Genes (DEGs) for RAP2.2 (GLYMA_19G262700) in each of the stress conditions, as determined by GENIE3 results and utilized their log2 fold change values for distance calculation. The resulting plots illustrate the similarities and differences between the various stress conditions based on this gene list. A distance of 0 signifies the minimum distance and thereby more similarity, while the maximum value represents the furthest distance and thereby more differences between the stress conditions. This approach provides a measure of dissimilarity between the selected gene lists differential expression profiles and helps identify patterns and relationships within the stress conditions.

### IMPRes *in silico* hypothesis generation

IMPRes (Integrative MultiOmics Pathway Resolution; Jiang et al., 2020a, 2020b), an innovative algorithm designed to address the challenges of extracting valuable insights and identifying active pathways by using complex multiomics data as evidence was used. RNA-seq data for control and various stress combinations and 6 input seed genes (Supplemental Data Set 14) which stood out from differential analysis, along with the soybean pathways background network generated using KGML files from KEGG were used. The generated results show a pruned network diagram (Figure 7A) showcasing the interconnections among the various active pathways that connect the seed genes and other genes in the pathways.

### Multiomics integrative correlation analyses

The mixOmics (Rohart et al., 2017) R package was used to correlate the transcriptomics data with reproductive (number of pods/plants, weight of pods/plant, pods with 1 seed, pods with 2 seeds, pods with 3 seeds, seeds/plant, seeds area, number of flowers/plant) and vegetative (growth rate, relative water content, Chlorophyll a, Chlorophyll b, total Chlorophyll, photosynthesis, transpiration, stomatal conductance, biomass) phenotypes. The transcriptomics data are the combination of DEGs identified with *q*-value less than 0.01 and log2 fold change (log2FC) of 2 for all comparisons. This cutoff extracted 5913 DEGs for the leaf group and 464 DEGs for the flower group. The data integration and classification were carried by Data Integration Analysis and Biomarker discovery using Latent variable approaches for Omics studies (DIABLO). We used the N-integration Sparse Patrial Least Square Discriminant Analysis (SPLS-DA) approach with function block.splsda to identify signatures composed of highly correlated variables across the multiple matrix sets, which enable us to detect a confident relationship between the data sets. The circos correlation matrix generated by the circosplot function was imported to the circlize (Gu et al., 2014) R package to create the circos plot. The gene expression correlated with vegetative and reproductive phenotypes within leaf group with a correlation cutoff of 0.8 and 0.92, respectively, while the flower group had a correlation cutoff of 0.7.

### Statistical analysis

For data shown in the main Figures, statistical analysis was performed using one-way ANOVA followed by Tukey’s *post hoc* test comparing the mean of each group with the mean of every other group in GraphPad. Different letters denote statistical significance at *P* < 0.05. Results are shown as box-and whisker plots with borders corresponding to the 25^th^ and 75^th^ percentiles of the data and the center line indicate the median. For data shown in Supplemental Figures data from each treatment was compared to control using Student’s *t*-test and asterisks indicate statistical significance at *P* < 0.05.

### Data availability

Transcript abundance and differentially expressed transcripts can be accessed interactively via the Differential Expression tool in SoyKB (https://soykb.org/DiffExp/diffExp.php; Joshi et al., 2012, 2014), a comprehensive all-inclusive web resource for soybean. RNA-Seq data was deposited in Gene Expression Omnibus (GEO), under the following accession number: GSE237798.

## ACKNOWLEDGMENTS

This work was supported by funding from the National Science Foundation (IOS-2110017, IOS-1353886, IOS-1932639), and Interdisciplinary Plant Group, and University of Missouri.

## AUTHOR CONTRIBUTION

M.A.P.V, R.S., S.P.I, Z.L., S.D.V., D.V., P.S., M.S.I., L.S.P. and B.S. performed experiments and analyzed the data. R.M., F.B.F, T.J., M.A.P.V, D.M.C, and S.I.Z. designed experiments, analyzed the data, and/or wrote the manuscript.

## Supplemental Data Legends

**Figure S1.**
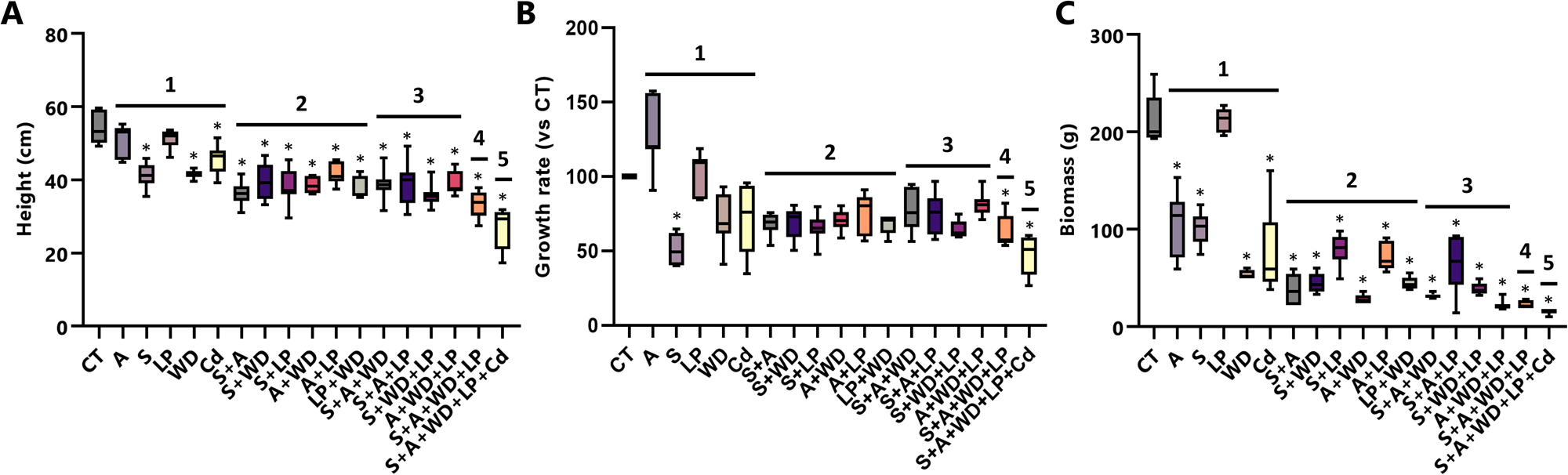
The effects of multifactorial stress combination (MFSC) on soybean height, growth rate and biomass (In support of Figure 1). **A.-C.** The effects of each stress/stress combination/MFSC on soybean height (A), growth rate (B), and biomass (C), respectively. Experiments were repeated in 3 biological repeats, each with 7 technical repeats. Results are shown as box-and whisker plots with borders corresponding to the 25^th^ and 75^th^ percentiles of the data. Statistical analysis was performed using Student’s t-test and asterisks denote statistical significance at *P* < 0.05. Abbreviations: A, acidity; Cd, cadmium; CT, control; LP, low phosphate; S, salinity; WD, water deficit.

**Figure S2.**
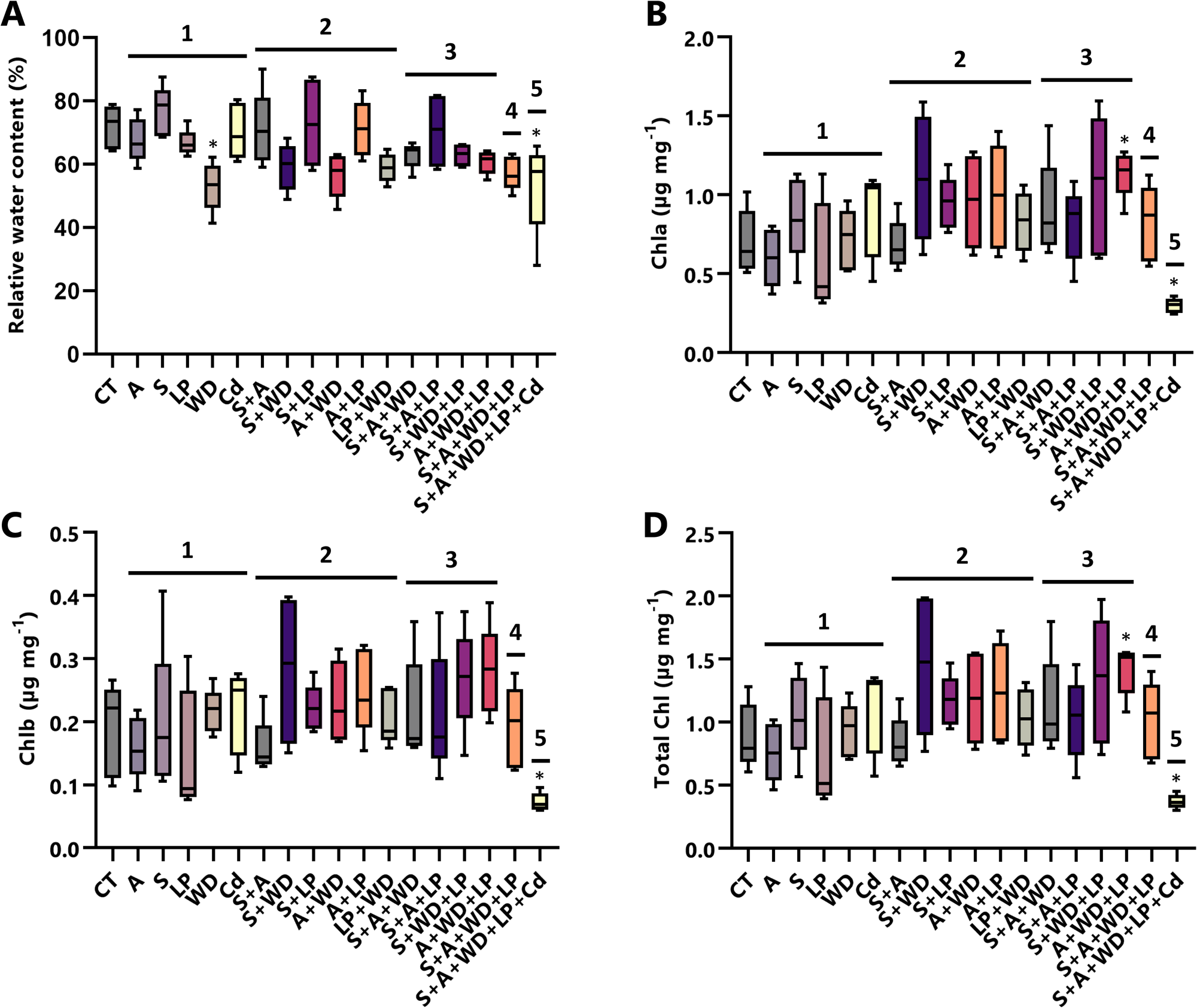
The effects of multifactorial stress combination (MFSC) on soybean relative water and chlorophyll content (In support of Figure 1). **A.-D.** The effects of each stress/stress combination/MFSC on soybean relative water content (A), and chlorophyll a (B), chlorophyll b (C), and total chlorophyl (D) content, respectively. Experiments were repeated in 3 biological repeats, each with 7 technical repeats. Results are shown as box-and whisker plots with borders corresponding to the 25^th^ and 75^th^ percentiles of the data. Statistical analysis was performed using Student’s t-test and asterisks denote statistical significance at *P* < 0.05. Abbreviations: A, acidity; Cd, cadmium; Chl, chlorophyll; CT, control; LP, low phosphate; S, salinity; WD, water deficit.

**Figure S3.**
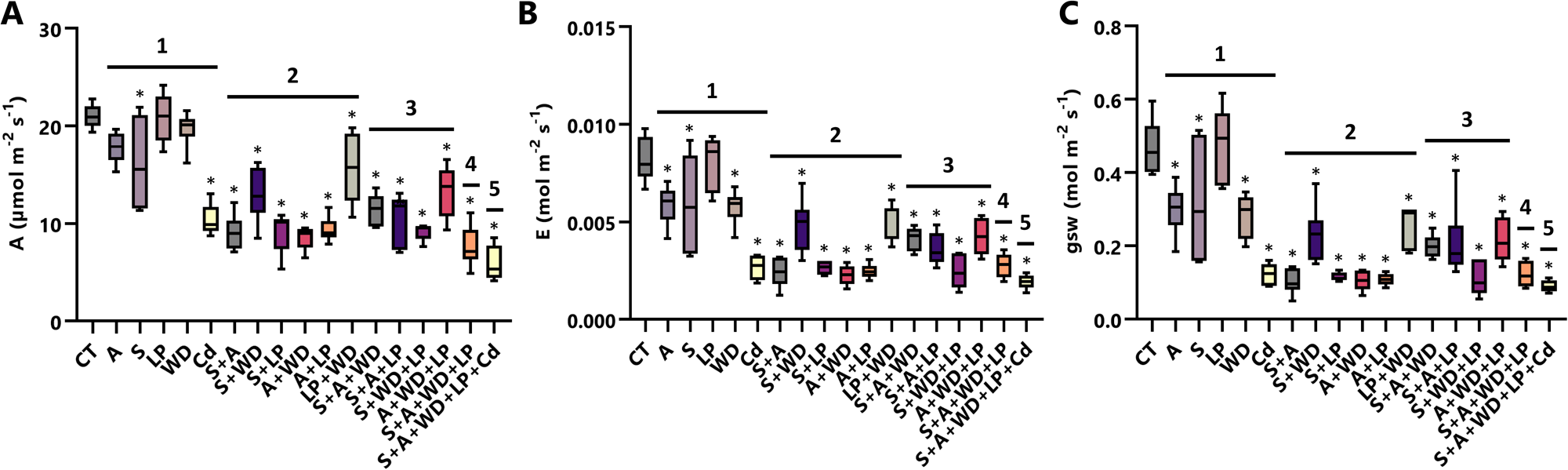
The effects of multifactorial stress combination (MFSC) on soybean physiological parameters (In support of Figure 2). **A.-C.** The effects of each stress/stress combination/MFSC on soybean leaf photosynthesis (A), transpiration (B), and stomatal conductance (C), respectively. Experiments were repeated in 3 biological repeats, each with 7 technical repeats. Results are shown as box-and whisker plots with borders corresponding to the 25^th^ and 75^th^ percentiles of the data. Statistical analysis was performed using Student’s t-test and asterisks denote statistical significance at *P* < 0.05. Abbreviations: A, acidity (X-axis); A, photosynthesis (Y-axis); Cd, cadmium; CT, control; E, transpiration; gsw, stomatal conductance; LP, low phosphate; S, salinity; WD, water deficit.

**Figure S4.**
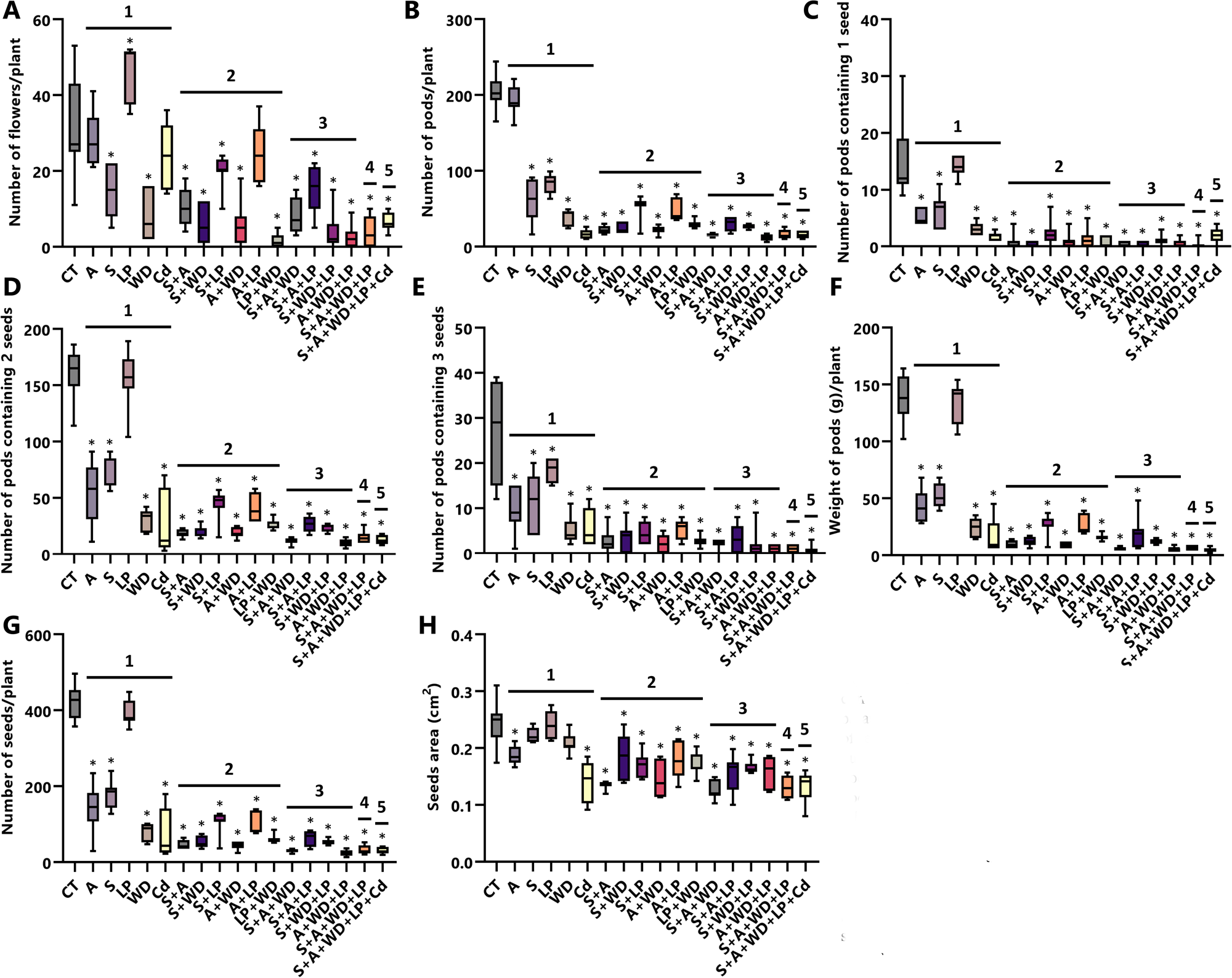
The effects of multifactorial stress combination (MFSC) on soybean reproductive parameters (In support of Figure 2). **A.-H.** The effects of each stress/stress combination/MFSC on soybean number of flowers (A), number of pods (B), number of pods containing 1 (C), 2 (D), or 3 (E) seeds, weight of pods (F), number of seeds per plant (G), and seed area (H), respectively. Experiments were repeated in 3 biological repeats, each with 7 technical repeats. Results are shown as box-and whisker plots with borders corresponding to the 25^th^ and 75^th^ percentiles of the data. Statistical analysis was performed using Student’s t-test and asterisks denote statistical significance at *P* < 0.05. Abbreviations: A, acidity; Cd, cadmium; CT, control; LP, low phosphate; S, salinity; WD, water deficit.

**Figure S5.**
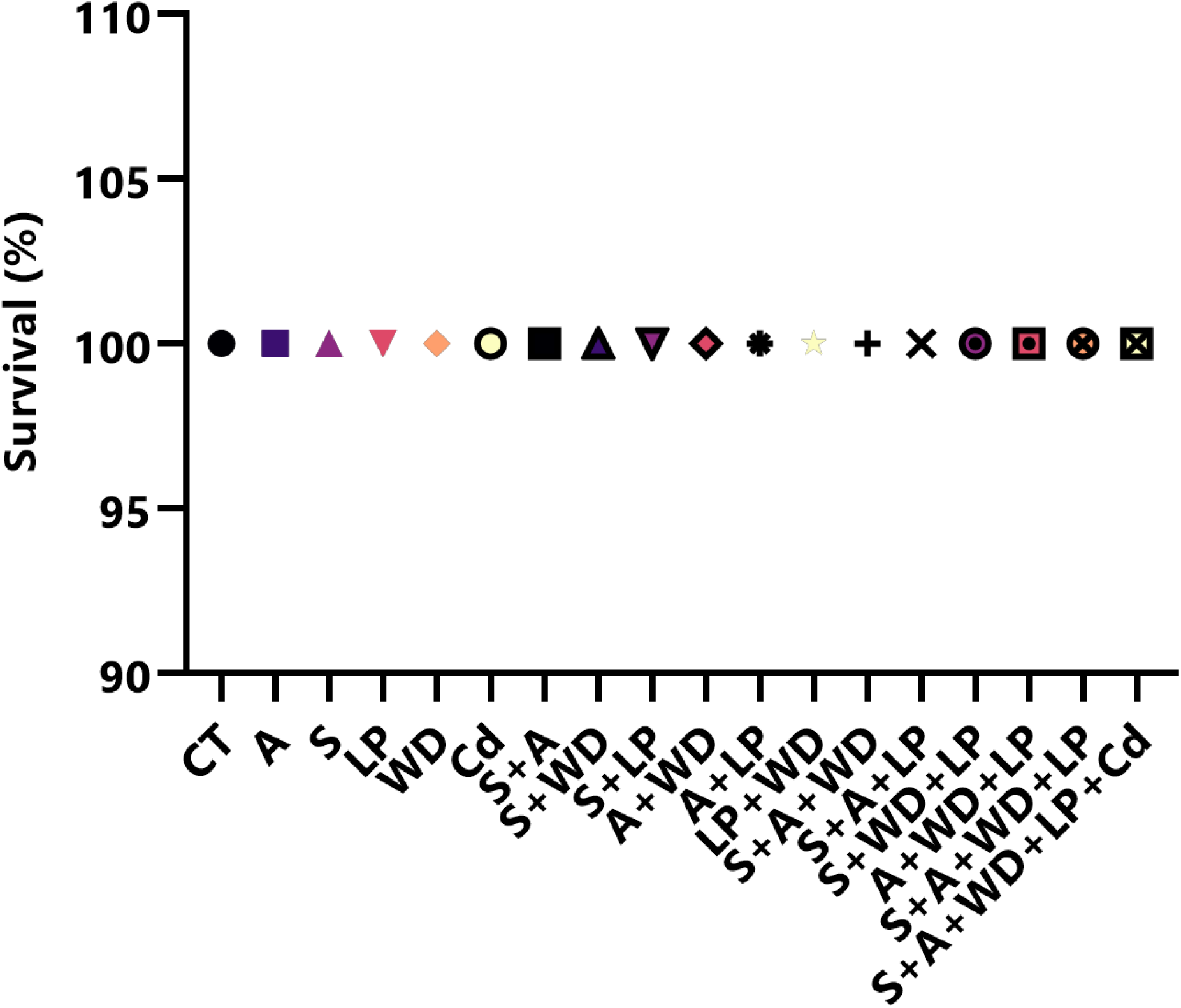
The effects of each stress/stress combination/MFSC on soybean seedlings survival (In support of Figures 1 and 2). Experiments were repeated in 3 biological repeats, each with 7 technical repeats. Results are shown as box-and whisker plots with borders corresponding to the 25^th^ and 75^th^ percentiles of the data. Statistical analysis was performed using Student’s t-test and asterisks denote statistical significance at *P* < 0.05. Abbreviations: A, acidity; Cd, cadmium; CT, control; LP, low phosphate; S, salinity; WD, water deficit.

**Figure S6.**
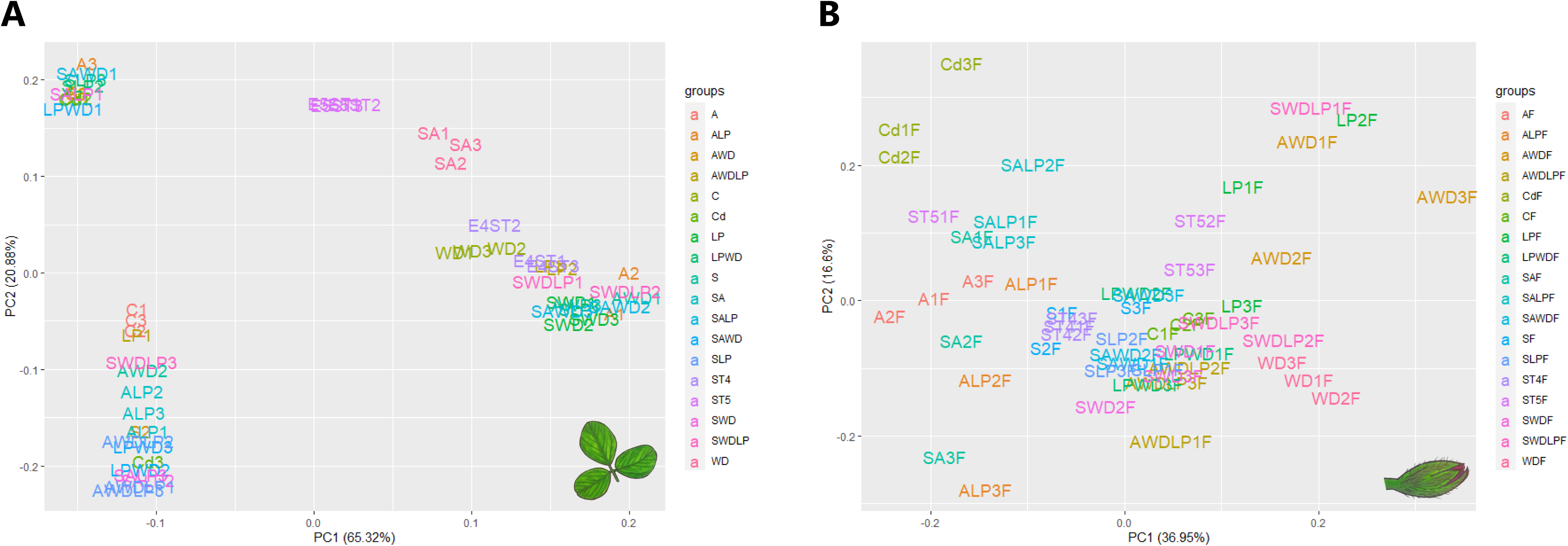
Principal component analysis (PCA) of transcriptomic data sets obtained from soybean leaf (A) or Flowers (B) subjected to each stress/stress combination/MFSC (In support of Figures 3-7). Experiments were repeated in 3 biological repeats, each with a pool of 7 technical repeats. Differentially expressed transcripts were defined as those that had an adjusted *P* < 0.05 (negative binomial Wald test followed by Benjamini–Hochberg correction). Abbreviations: A, acidity; CT, control; LP, low phosphate; S, salinity; WD, water deficit.

**Supplemental Data Set 1.** Transcripts differentially expressed in leaves of soybean plants subjected to water deficit (WD), acidity (A), low phosphate (LP), salinity (S) and cadmium (Cd) stresses alone and in combination including 1, 2, 3, 4 and 5 stresses.

**Supplemental Data Set 2.** Transcripts differentially expressed in flowers of soybean plants subjected to water deficit (WD), acidity (A), low phosphate (LP), salinity (S) and cadmium (Cd) stresses alone and in combination including 1, 2, 3, 4 and 5 stresses.

**Supplemental Data Set 3.** Overlap between transcripts significantly altered in soybean leaves in response to all 1-, 2-, 3- and 4-stress combinations (Figure 3A).

**Supplemental Data Set 4.** Overlap between transcripts significantly altered in soybean flowers in response to all 1-, 2-, 3- and 4-stress combinations (Figure 3B).

**Supplemental Data Set 5.** Transcripts common to the response of soybean leaves and flowers to all 1-, 2- and 3-stresses combination (Figure 4A) and 1+2+3 and 4-stress combinations (Figure 4B) and their overlap.

**Supplemental Data Set 6.** Gene ontology and KEGG pathway enrichment of the transcripts identified in the soybean leaf and flower transcriptomic response to all 3- and 4-stress combinations.

**Supplemental Data Set 7.** Overlap between transcripts significantly altered in soybean leaves and flowers in response to all 5 single stresses (Figure 5A). Overlap between transcripts significantly altered in soybean leaves and flowers in response to all 3-, 4-, and 5-stress combinations (Figure 5B). Venn diagram comparing the response of soybean leaves and flowers to cadmium and to the 5-stress combination (Figure 5C).

**Supplemental Data Set 8.** Gene ontology and KEGG pathway enrichment of the transcripts identified in soybean leaves and flowers transcriptomic responses to all 3-, 4- and 5-stress combinations.

**Supplemental Data Set 9.** Comparison between all transcripts significantly altered in Arabidopsis seedlings and soybean leaves in response to MFSC (Figure 5D). Gene ontology and KEGG annotation analysis of the transcripts common to the response of Arabidopsis and soybean to multifactorial stress combination (Figure 5E).

**Supplemental Data Set 10.** GENIE3 network analysis results for HSFA8, HSFA4A, HSFA3, HSFA4A, RAP2.4, RAP2.5 and RAP2.2 transcription factors in soybean leaves.

**Supplemental Data Set 11.** GENIE3 network analysis results for HSFA8, HSFA4A, HSFA3, HSFA4A, RAP2.4, RAP2.5 and RAP2.2 transcription factors in soybean flowers.

**Supplemental Data Set 12.** List of transcripts associated with RAP2.2 and HSFA4A transcription factors in leaves of soybean plants subjected to S+A+WD+LP+Cd stress (Figure 6C top).

**Supplemental Data Set 13.** List of transcripts associated with HSFA8 transcription factor in flowers of soybean plants subjected to S+A+WD+LP+Cd stress (Figure 6C bottom).

**Supplemental Data Set 14.** IMPRes *in silico* pathway analysis of the response of soybean flowers to MFSC (Figure 7A).

**Supplemental Data Set 15.** MixOmics analysis linking specific transcripts expressed in leaves or flowers, with specific vegetative or reproductive phenotypes (Figure 7B-E).

